# The cellular basis of feeding-dependent body size plasticity in sea anemones

**DOI:** 10.1101/2023.05.15.540851

**Authors:** Kathrin Garschall, Eudald Pascal-Carreras, Belén Garcia-Pascual, Daria Filimonova, Annika Guse, Iain G. Johnston, Patrick R.H Steinmetz

## Abstract

Animals with indeterminate growth can adapt their growth rate and body size to changing food availability throughout their lifetime. As the cellular basis underlying food-dependent growth plasticity is poorly understood, we quantified how the sea anemones *Nematostella vectensis* and *Exaiptasia diaphana* (Aiptasia) respond to feeding and starvation on organismal and cellular levels. Using mathematical modelling to analyse growth phases, we found that growth and shrinkage rates in *Nematostella* are exponential, stereotypic and accompanied by high levels of cell gain or loss, respectively. During starvation and re-feeding, a considerable proportion of juvenile polyp cells (>7%) reversibly shift between S/G_2_/M and G_1_/G_0_ cell cycle phases, suggesting a tight nutritional control of quiescence and cell cycle re-entry. In the facultative symbiotic sea anemone Aiptasia, we found that growth and cell proliferation rates are dependent on the symbiotic state and, in comparison to *Nematostella*, respond less strongly to changes in food supply. Altogether, we provide a benchmark and resource to study the nutritional regulation of body plasticity on molecular, cellular and genomic levels using the rich functional toolkit available for *Nematostella*.

**Summary statement:** Feeding and starvation in sea anemones induce growth and shrinkage, cell size changes and dynamic cell proliferation changes that support a nutritional control of quiescence and cell cycle re-entry.

## Introduction

Indeterminate growth, the ability to control growth in response to environmental conditions, is likely ancestral to animals and found among many bilaterians (e.g., fish, planarians or crustaceans) and non-bilaterians (e.g., cnidarians, ctenophores, sponges) (Hariharan et al., 2015; Sebens, 1987). The ability to plastically adapt growth rates and body size to food availability often comes with a high resilience to starvation stress (McCue, 2010). Whole-body shrinkage is an extreme starvation response common in ctenophores (Jaspers et al., 2015), cnidarians (Lilley et al., 2014), acoels (De Mulder et al., 2009; Nimeth et al., 2004) and planarians (Baguñà J. and Garcia-Fernandez, 1990), and sporadically found in vertebrates (lampreys (Olesen Larsen, 1962), eels (Boëtius and Boétius, 1985), amphibians (Bendik and Gluesenkamp, 2013), reptiles (Field et al., 2007; Wikelski and Thom, 2000)) and some protostomes (e.g., polychaetes (Åkesson and Rice, 1992) or gastropods (Böer et al., 2007)).

Cnidarians (e.g., sea anemones, corals, jellyfish or hydroids) exhibit high body plasticity and indeterminate growth (Sebens, 1987; Technau et al., 2015). Feeding-dependent growth and starvation-induced whole-body shrinkage are observed in scyphozoans (’true jellyfish’) (Frandsen and Riisgård, 1997; Hamner and Jenssen, 1974; Lilley et al., 2014), hydrozoans (e.g. *Hydra*, *Clytia*) (Loomis, 1953; Matsakis, 1993; Muscatine and Lenhoff, 1965; Otto and Campbell, 1977)) and anthozoans (sea anemones and corals) (Chomsky et al., 2004; Sebens, 1980)).

Currently, little is known about the cellular or molecular processes underlying the feeding-dependent adaptation of growth rates in animals. A main reason is that the evolutionary derived, genetic pre-determination of body sizes found in most genetic research organisms (e.g. mammals, insects, nematodes) leads to an uncoupling of feeding and growth rates during maturation (Hariharan et al., 2015). How feeding regulates body growth is best studied on a cellular level in the hydrozoan *Hydra* (Bosch and David, 1984; Buzgariu et al., 2014) and the planarian flatworm *Schmidtea mediterranea* (Baguñà J. and Garcia-Fernandez, 1990; González-Estévez et al., 2012; Newmark and Sanchez Alvarado, 2000). Despite their phylogenetic unrelatedness, adult stem cells in *Hydra* and planarians (i.e., neoblasts) sustain a high overall cell turnover, and respond similarly to feeding and starvation (Baguñà and Romero, 1981; Baguñà J. and Garcia-Fernandez, 1990; Otto and Campbell, 1977). In both animals, feeding induces a cell proliferation burst that fades back to a baseline level staying stable even during long starvation periods. Body shrinkage in both *Hydra* and planarians results from increased levels of apoptosis and changes in cell cycle dynamics during starvation leading to net cell loss (Baguñà and Romero, 1981; Bosch and David, 1984; Böttger and Alexandrova, 2007; González-Estévez et al., 2012). Cell cycles of *Hydra* stem cells are unusual with a nearly absent G_1_ phase and a long G_2_ phase (Campbell and David, 1974; David and Campbell, 1972; Holstein et al., 1991). Starvation extends their cell cycle lengths up to 2-fold but does not induce an arrest, which occurs in in G_2_ during cell differentiation (Bosch and David, 1984; Dübel and Schaller, 1990). Planarians appear to have more traditional cell cycle phases, but it is currently unclear if starvation leads to their cell cycle to slow-down or completely arrest (Kang and Alvarado, 2009; Molinaro et al., 2021; Newmark and Alvarado, 2000).

While planarians and *Hydra* have provided valuable cellular insights into the nutritional regulation of growth and body size, the universality of their strategies is difficult to assess given the limited sampling of taxa and metabolic behaviours. To this end, we here reveal the quantitative organismal and cellular dynamics of indeterminate growth in two sea anemones with and without phototrophic symbionts. Specifically, we focussed on the sea anemones *Nematostella vectensis* and *Exaiptasia diaphana* (Aiptasia), the latter being facultatively associated with symbiotic dinoflagellate algae that provide photosynthetically produced carbohydrates to the host (Davy et al., 2012; Rädecker et al., 2018). In *Nematostella*, feeding affects growth (Fig. 1A) (Carvalho et al., 2023; Hand and Uhlinger, 1992) and is necessary for the induction of tentacle growth (Ikmi et al., 2020). During starvation, *Nematostella* polyps shrink (Fig. 1A), reduce proliferation, retain characteristic locomotory behaviour and rescale the number of GLWamide neuropeptide-expressing neurons proportionally to their reduced size (Havrilak et al., 2021; Passamaneck and Martindale, 2012). In the facultative symbiotic Aiptasia, growth is feeding-dependent but it is unclear if symbionts provide additional benefits to support growth or buffer shrinkage rates (Clayton Jr and Lasker, 1985; Leal et al., 2012; Tivey et al., 2020). Until now, the quantitative dynamics of feeding- or starvation-induced changes in growth, shrinkage, cell size, cell numbers and proliferation rates remain poorly understood in these species or any other animals, except *Hydra* and planarians.

**Fig. 1.**
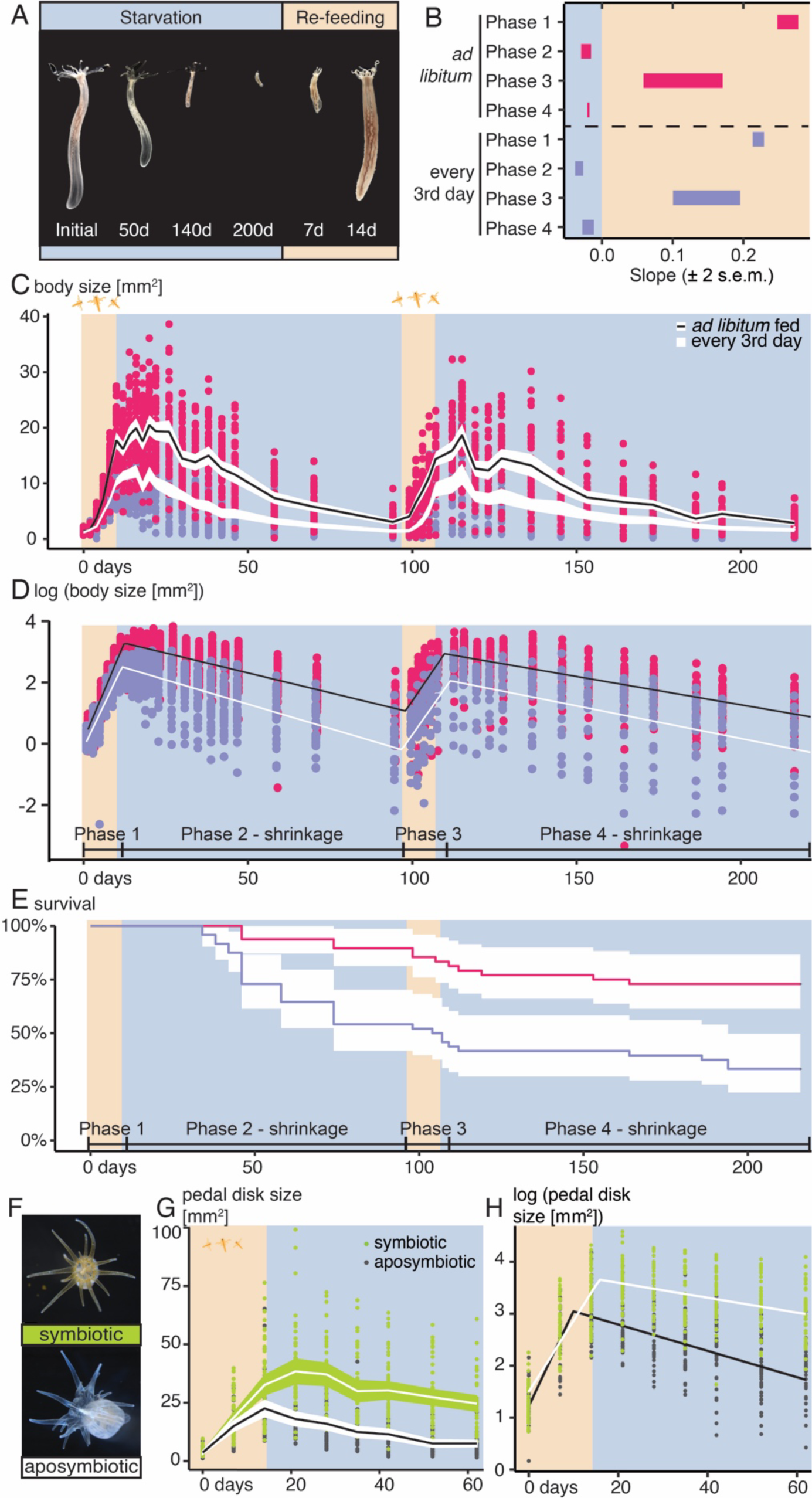
Feeding-dependent body size plasticity in *Nematostella* (A-E) and Aiptasia (F-H). **(A, F)** Representative images of *Nematostella* juveniles during starvation and refeeding (A) and of symbiotic and aposymbiotic Aiptasia strain CC7 juveniles **(F)**. **(B-E)** Comparison of growth and shrinkage rates **(B)** and quantification of body size **(B, G)**, log-transformed body size **(C, H)** and survival rates **(E)**. **(B)** Estimates of growth and shrinkage rates from fitting linear models to log-transformed body size values from *ad libitum* (pink) and polyps fed every 3rd day (purple). **(C-D)** Body size **(C)** and survival rates **(E)** of juvenile *Nematostella* polyps was measured over two 10d-feeding-90d-starvation cycles for *ad libitum* fed or restricted feeding regimes. Individual values (dots) and mean values with a 95% confidence interval are plotted and connected across different timepoints. n=96 per condition. **(D)** A linear model with changepoints fitted to the log-transformed values of body area. Traces illustrate mean behavior of the 4-phase model favored by model selection (see SI and Fig.S1) for black (*ad libitum*) or white (every 3^rd^ day) lines. (**E)** Kaplan-Meier plot of survivorship for the feeding regimes. Cox proportional hazard showed significantly reduced survival in the restricted fed animals (Hazard Ratio = 3.61, 95% confidence interval (1.89, 6.90), p < 0.001). **(G)** Pedal disk area was measured during 2 weeks of feeding and 8 weeks of starvation; n=36 animals per condition. **(H)** As in (D) of Aiptasia pedal disk area during feeding and starvation. Traces illustrate mean behaviour of the 2-phase model favored by model selection (see Fig.S5 and SI).

## Results and discussion

### Feeding-dependent body size plasticity in *Nematostella*

We tested the effect of different feeding regimes on body growth rates in *Nematostella* by measuring changes in body column area of *ad libitum* or restricted fed (every 3rd day) juveniles (Reitzel et al., 2013) over two consecutive 10 days-fed/90 days-starved cycles (Fig. 1B-D). We observed that fed animals grew and shrank exponentially at feeding-dependent rates, which we captured quantitatively by exploring a range of changepoint-coupled linear models for log-transformed size with time (Fig. 1B-D, Fig. S1). Changes in body size were best described by a four-phase model: two growth phases (Fig.1D, phases 1 and 3) and two shrinkage phases (Fig. 1D, phases 2 and 4). These phases largely overlapped with the feeding/starvation intervals, but the growth phases extended a few days into the starvation intervals, before shrinkage started, for all feeding regimes (Fig. 1D, Supplementary Information Table 5 for changepoints).

During the first growth phase, we found major feeding-dependent differences between growth rates of *ad libitum* (∼13-fold increase, doubling time T_D_: 2.5–3.0 days) and restricted feeding (∼9-fold increase, T_D_: 2.9–3.2 days) (Fig. 1B,D, SI Table 5,8). The size difference between the regimes that resulted from these initial growth rates remained throughout the entire experiment (Fig.1B). During the second feeding phase, polyps under both feeding conditions generally grew slower during the initial phase (Fig. 1D; compare phase 3 to phase 1). Surprisingly, polyps under restricted feeding tended to grow faster (T_D_: 3.5–5.9 days) than continuously fed animals (T_D_: 4.2–6.1 days) during phase 3 (Fig. 1D, SI Table 8), possibly due to compensatory growth from a smaller starting size after the starvation phase 2. We also asked how the different dimensions of the polyp body column change during growth, and found that body length strongly correlates to width^2^, meaning that animals grow predominantly in length over ten days of feeding (R^2^: 82%, Fig. S2E,F, Table S1).

Shrinkage rates strongly differed between feeding regimes during the shrinkage phases 2 and 4 (Fig. 1D, SI table 4 and table 9). During phase 2, *ad libitum* fed animals halved their size every 4 weeks (halving time T_1/2_: 25.8–30.4 days), while polyps with restricted feeding shrank much more quickly (T_1/2_: 18.6–23.6). During phase 4, the difference between feeding regimes was less pronounced (T_1/2_ (ad libitum): 34.3–41.1 days; T_1/2_ (restricted): 27.4–40.5 days). The generally lower shrinkage rates in both feeding groups point to potentially increased starvation tolerance during phase 3. By comparing survival rates between feeding regimes (Fig. 1E), we found that lethality was higher under restricted feeding during shrinkage phase 2. During phase 4, however, lethality was generally lower and similar between feeding regimes. This could be explained by selection for polyps with higher starvation resistance during phase 2. Interestingly, lethality also increased during refeeding in both groups (Fig. 1E, phase 3), suggesting that metabolic changes accompanying the starving-to-feeding transition impose organismal stress.

To estimate the stereotypy of growth between batches and experiments, we compared growth rates between equivalent but independent experiments (Fig. S2A-D, SI Table 1 and Table 4). For all growth experiments, we found that simple, single-phase linear regression models were a good fit to the data and tested if the 95% confidence interval of their slopes intersected (Fig. S2D). We observed that confidence intervals on the slopes of *ad libitum* growth phases largely overlapped when not preceded by starvation (Fig. S2D, ‘0-10d fed’ & ‘*a.l.* phase 1’). In addition, confidence intervals for growth rates also matched between all samples refed after starving between 90 and 200 days (Fig. S2D, all ‘phase 3’ and ‘refed’ samples). Together, and given the reasonably tight bounds on estimate slopes reflected by these intervals, this suggests a stereotypic and reproducible growth response to feeding depending on starvation history.

Body size changes during starvation were analysed using linear modelling with changepoints delimiting dynamic “phases” with different slopes, determining the best fitting model via Akaike Information Criterion (AIC) for up to eight different phases (Fig. S3A-G, see Methods and Supplementary Information). In the three-phase scenario (200 days starvation, Fig. S3C-G), an initial, short exponential growth phase was followed by an exponential shrinkage phase and a relatively flat plateau phase after >100 starvation days. Inference of the slopes in each phase and comparing their 95% confidence intervals showed that exponential shrinkage phases are similar between experiments (Fig. S3H). A simpler approach, performing linear regression on manually isolated intervals between peaking body size and >100 days of starvation resulted in similar slopes with halving rates ranging between 21.6 and 35.4 days for all starvation experiments (Table S1). Together, our data suggests that animals grow about 7-10 times faster than they shrink. The general consistency of shrinkage rates between experiments shows that the response to starvation in juvenile *Nematostella* is stereotypic and highly reproducible.

We then tested if the growth and starvation responses are comparable between the estuarine sea anemone *Nematostella* and the tropical marine sea anemone Aiptasia. Aiptasia, which can be studied in the presence or absence of photosynthetic algae, is of particular interest in assessing the effect of symbionts on growth and shrinkage dynamics (Fig. 1F). We found that *ad libitum* fed Aiptasia polyps show similar growth rates (≈ changes in pedal disk area (Leal et al., 2012)) between symbiotic (T_D:_ 2.9– 5.2 days) and aposymbiotic animals (T_D_: 3.1–4.9 days, Fig. 1G,H Fig. S5G, SI Table 11). Symbiotic polyps showed continuous growth during the first days of starvation. In contrast, aposymbiotic polyps started shrinking immediately upon starvation at rates that were higher (T_1/2_: 23.9–33.0 days) than in symbiotic polyps (T_1/2_: 46.2–151.3 days) over 8 weeks (Fig.1 G,H). Generally, Aiptasia grew about 3-7x faster than they shrunk (SI Table 12,13. Altogether, Aiptasia and *Nematostella* responded roughly similarly to feeding and starvation. Under the given temperature conditions, *Nematostella* showed generally higher shrinkage rates than Aiptasia. Experiments investigating the combined effect of food and temperature will be necessary to estimate if shrinkage rates are generally higher in *Nematostella* than in Aiptasia, for example due to higher metabolic rates or lower nutrient storage capacities.

### Cell size and numbers strongly correlate with nutrient supply in *Nematostella*

To test how cell numbers and/or cell size change during growth or shrinkage in *Nematostella*, we established an imaging and flow cytometry workflow to quantify body size, global cell numbers, median cell size and cell cycle distribution of individual polyps. Quantification of total cell numbers was both sensitive and scalable over a wide range of cell numbers in *Nematostella* juveniles (Fig. S6A). Over 10 days of feeding, cell numbers increased exponentially (Fig. 2C, T_D_∼3.2 days) and log cell number correlated well with log body size changes (Fig. 2E, R^2^: 65%). Notably, median cell sizes increased over 10 days of feeding (+46%, Fig. 2D) without appearing to reach a maximum, showing that larger polyps had on average larger cells (Fig. 2F; log space R^2^: 48%). This observation could result from a shift towards larger cell types during polyp growth and/or a global increase in cell size in larger and well-fed animals. A linear regression model in log-transformed space provided a reasonable fit between both cell number and cell size and body size, suggesting that a power law relationship of the approximate form (body size) ∼ (cell size)^1^(cell number)^2/3^ links cell and body properties during growth (SI Geometry, R^2^: 68%).

**Fig.2.**
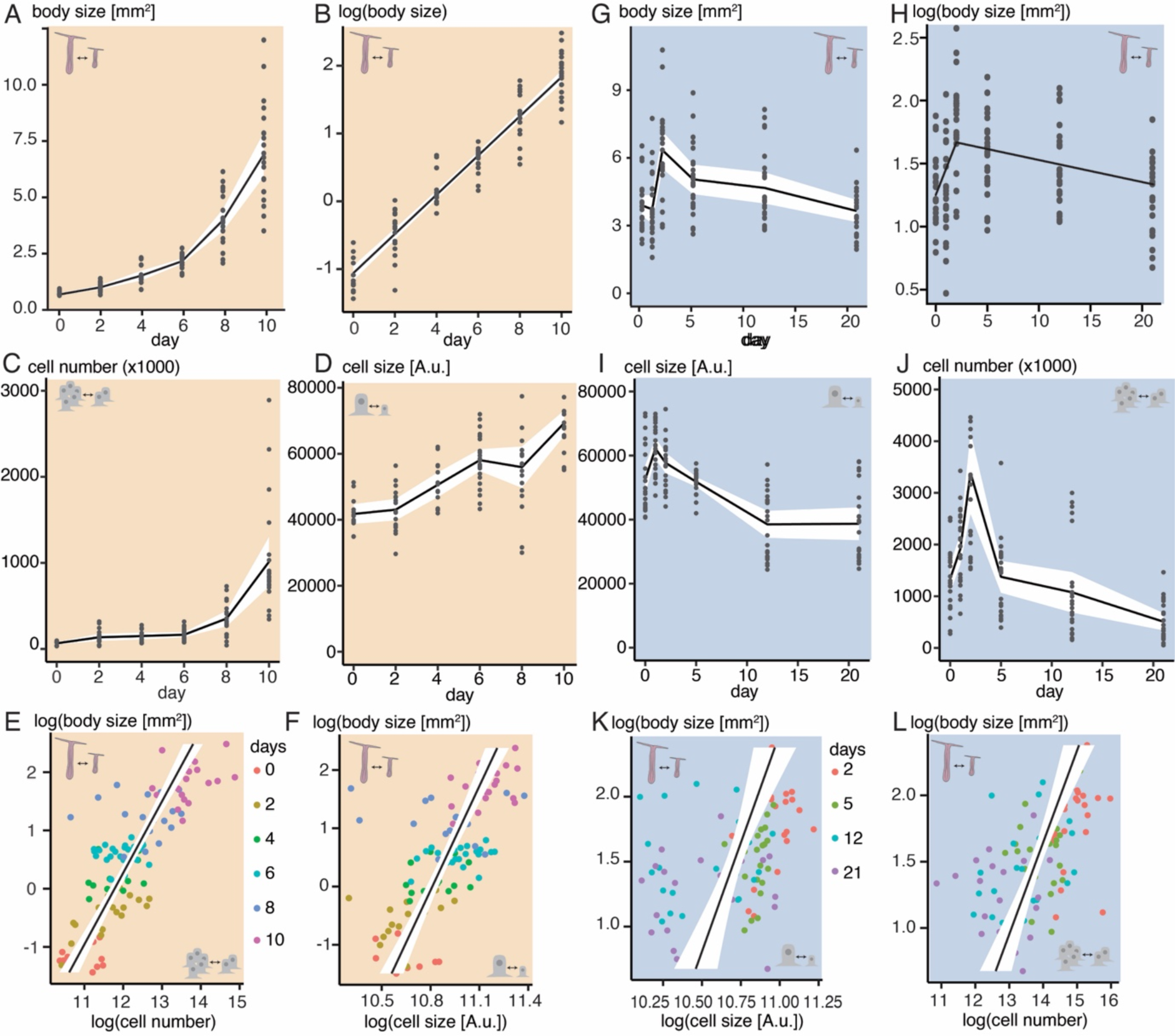
Body size, cell number and cell size change during feeding and starvation. Body size **(A,B,G,H,E-L)**, cell number **(C,J,E,L)** and cell size **(D,I,F,K)** of individual juvenile polyps were measured during 10-days of *ad libitum* feeding **(A-F)** and 21 days of starvation **(G-L)**. **(A, G, C, D, I, J)** show means and 95% confidence intervals connected across timepoints. **(B)** shows linear model fit to log-transformed data. **(E, F, K, L)** Correlations between log body size and cell number (E, L) and cell size **(F, K)**. **(H)** shows the best fit changepoint model on the log-transformed data (see Fig.S4A, SI).For all: n=24 per timepoint.

Median cell sizes peaked after 1 day of starvation (+18%) before decreasing and reaching a plateau after 12 days (−38% since T_0_) (Fig. 2I, Fig. S4E-H). In contrast to fed animals (Fig. 2F), the relationship between log cell size and log body size was too weak in starved animals to provide significant predictive power (R^2^: 15%, Fig. 2K, SI Table 7), suggesting that cell size may reflect feeding status rather than polyp size.

At the onset of starvation, body size and cell numbers initially increase over two days (Fig. 2G,J). Strikingly, this short growth phase is followed by a major cell loss over the next three (>50% loss) and 21 days (>80% loss)(Fig. 2J). Using a Neubauer chamber, we confirm a similar ∼86% reduction in cell numbers between 2 and 21 days post-feeding (Fig. S6B, Table S1). This dramatic cell loss was surprising as the median body size decreased by only ∼45% during the same interval (Fig. 2G, Table S1). Consequently, log body size and log cell numbers only poorly correlated between 2 and 21 days of starvation, suggesting a weaker relationship than during the growth phase (R^2^: 30%, Fig. 2L, SI Table 7). This discrepancy could result from a disproportionately high cell loss in tissues excluded from our measurements (e.g., tentacles, mesenteries) and/or from a delay in extracellular matrix scaffold breakdown after cell loss. Future studies will be needed to clarify the relevance of these processes during starvation-induced tissue remodelling.

### Cell proliferation is strongly regulated by feeding in sea anemones

These cell number dynamics during growth suggest that nutrient supply regulates cell proliferation rates. We next tested this assumption by quantifying changes in the fraction of S-phase cells comparing confocal imaging-based and flow cytometry-based methods during feeding and starvation (Fig.3 A-G, Table S1). Using flow cytometry to detect cells labelled by the S-phase marker 5-ethynyl-2’-deoxyuridine (EdU), we found that the ratio of EdU+/EdU– cells (i.e., EdU index) dropped from 7.2±0.4% to 1,4±0.2% between 24h and 5 days post-feeding (’dpf’, Fig. 3G, Table S1). A minor, significant increase between 5 and 21 days post-feeding (to ∼2%; p = 0.0421, paired t-test) may suggest a preferential retention of proliferative cells during starvation (further tested below). Re-feeding at 21dpf led to a dramatic restoration of the EdU index to ∼8% within 24 hours (Fig. 3G, Table S1). Confocal imaging-based quantification showed higher variability, but broadly confirmed the EdU index changes obtained by flow cytometry (Fig. 3A-G, Table S1). The exponential cell number increase observed during growth is thus well explained by the remarkably high fraction of S-phase cells in fed animals. In addition, this S-phase fraction rapidly shrinks within 2 days of starvation, suggesting a tight regulation of cell proliferation by food availability in *Nematostella*.

**Fig.3.**
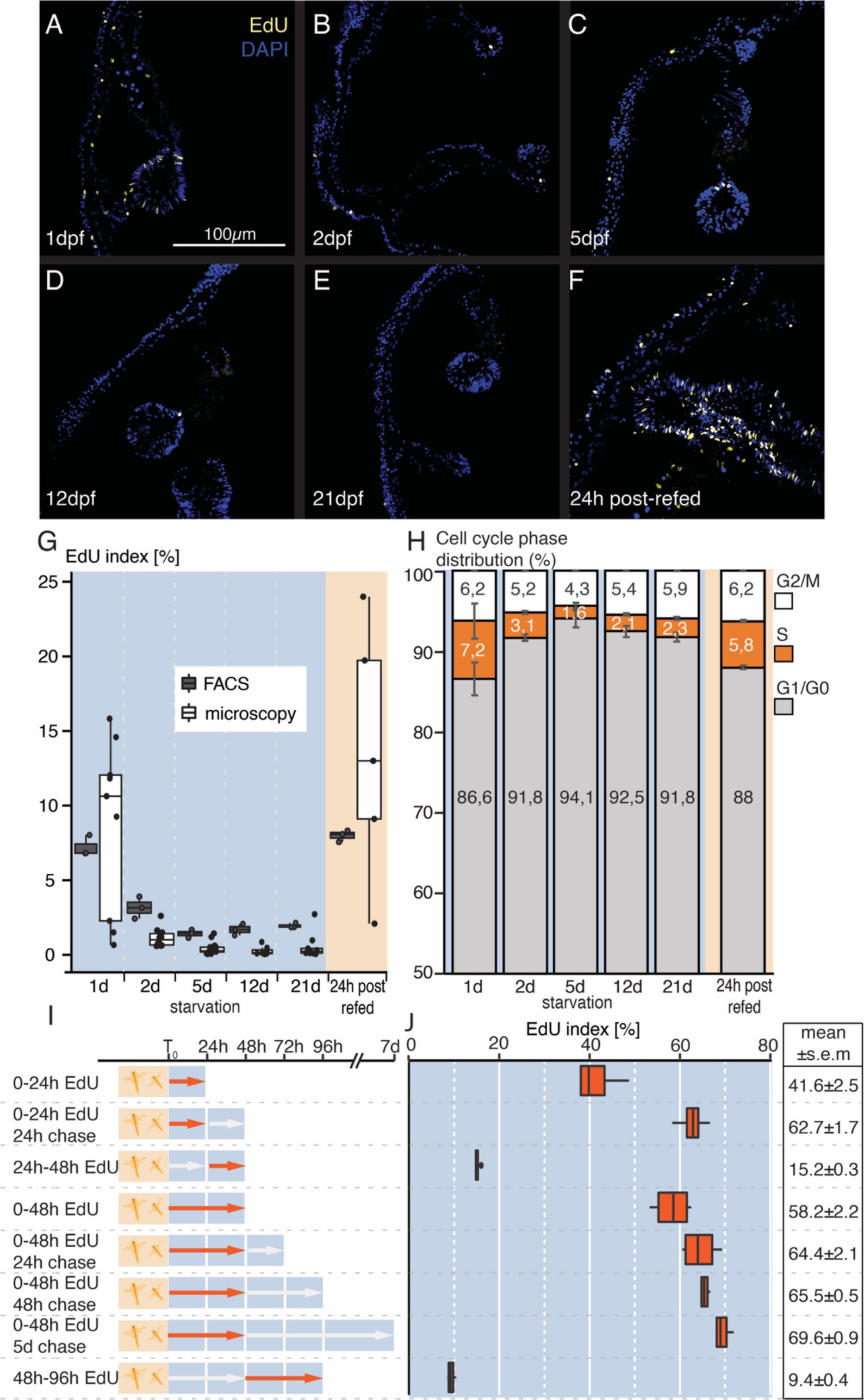
Feeding and starvation strongly affect cell proliferation and cell cycle phase distribution. Representative confocal images of EdU-labelled cells (A-F) and quantification of EdU index (G), cell cycle phase distribution (H) and EdU pulse-experiments by imaging (G) or flow cytometry (G-J). **(A)** Stacks of confocal images of juvenile mid-sections after 60min EdU labeling. Blue: DAPI nuclear stain. Yellow: EdU label. dpf: days post-feeding. h: hours. **(G)** Quantification of the fraction of EdU-positive cells by imaging analysis (white bars) and flow cytometry analysis (grey bars). n(FACS)= 3 biological replicates; n(microscopy)=5-14. **(H)** Cell cycle composition determined by flow cytometry based on DNA staining signal intensity. n=3 biological replicates **(I,J)** Experimental setup (I) and quantification of EdU+ fraction of cells in long pulse and chase experiments. n=4 biological replicates. Cell cycle composition and median fluorescence intensity are found in the supplementary Fig. S7A,B.

In Aiptasia, using a similar flow cytometry/EdU-based approach, we observed a significantly higher EdU index in symbiotic than in aposymbiotic polyps (Fig. S7D), supporting the higher growth rates observed in symbiotic polyps (Fig. 1G,H). Our results thus confirm and extend recent observations that the density of proliferating cells in Aiptasia tentacles increases near symbiotic cells and is generally higher than in aposymbiotic tentacles (Tivey et al., 2020). Compared to *Nematostella*, the Aiptasia EdU index ranged between similar values (between approx. 4.8 to 7.2%) and its starvation-induced decrease was less steep, independently of the symbiotic status. We thus conclude that the induction of high levels of cell proliferation as response to feeding is common among sea anemones (Actinaria).

### Starvation induces G_1_/G_0_ accumulation in a large pool of proliferating cells

The EdU/S-phase index provides a snapshot that only poorly reflects the total proportion of proliferating cells and provides no information about cell cycle phase dynamics. We therefore measured DNA content by flow cytometry to reveal changes in the proportion of cells in G_1_ (2N DNA content), S (between 2N and 4N) and G_2_/M (4N) phases (Fig. 3H). By quantifying the proportion of combined S/G_2_/M phases, we determined that at least 11,4%±0,7% of all cells are proliferative under *ad libitum* feeding in *Nematostella* juveniles (Fig. S7D). We furthermore tested if cells in starved animals predominantly accumulate in a specific cell cycle phase by comparing changes in G_1_, S and G_2_/M phase fractions (Fig. 3H, Table S1). The fraction of S-phase cells decreases between 24 and 48hpf from ∼7% to ∼3%, corresponding to an increase of the G_1_ fraction (from ∼87% to ∼92%) with a relative stable proportion of cells in G_2_ (from ∼6% to ∼5%) (Fig. 3H). Refeeding induced an increase of S-phase cells (to ∼6%) and a complementary reduction of G_1_ phase cells (to ∼88%) within 24 hours (Fig. 3H. Our data thus strongly suggest that starvation induces quiescence predominantly from G_1_ in a large proportion of proliferating cells, and that re-feeding induces their cell cycle re-entry. This remarkable capacity to arrest proliferative cells upon starvation and keep them competent to re-enter the cell cycle is likely underlying the nutritional control of growth, a key trait of indeterminately growing animals. Notably, this feature is absent in *Hydra* (Bosch and David, 1984) and likely not predominant in planarians (Newmark and Alvarado, 2000). Quiescent stem or progenitor cells are also extremely rare or absent in adults of genetic research animals (e.g., mammals, flies, nematodes), where growth is decoupled from nutrient supply.

### Long term EdU pulse substantiates high rates of cell addition at the onset of starvation

Using continuous EdU pulses, we aimed to estimate the proportion and fate of newly born cells during the first seven days of starvation (Fig. 3I). We found that 41.6±2.5% of all cells incorporate EdU between 0 and 24hpf (Fig. 3I,J). As >60% of these EdU+ cells were in G_1_ (Fig. S7C), a larger proportion of EdU+ cells underwent at least one recent mitosis and likely differentiated in G_1_. These observations broadly support the increase in cell number underlying body growth during the same period (Fig. 2E, Fig. S4D,L). A 24h pulse-24h chase experiment EdU-labelled 62.7±1.7% of all cells (Fig. 3I,J) but resulted in lower median EdU labelling intensities than after a 24h pulse/0h chase (Fig. S7B). This indicates that dilution of the EdU label occurred by continuing cell divisions between 24 and 48hpf, albeit at lower rates than during the first 24 hours. Continuous labelling between 24 and 48hpf results in a 15.2±0.3% EdU index (Fig. 3I) and confirms that cell proliferation activity continues but decreases during this starvation interval. Comparing the S/G_2_/M fraction of EdU+ cells between 0-24hpf (29.9±1.6%) and 24-48hpf (37.0±1.1%), this fraction of actively proliferating cells is significantly higher during the later interval (Fig. S7C, Table S1). This observation indicates a lengthening of the cell cycle and/or a decrease in the proportion of newly differentiating cells among all EdU+ cells. Correspondingly, the EdU index after a continuous 0-48hpf pulse (58.2±2.2%) corresponds roughly to the sum of the EdU indexes between 0-24h (41.6±2.5%) and 24-48h (15.2±0.3%) (Fig. 3J, Table S1). Given the dramatic drop in cell numbers between 2 and 5 dpf, we were interested to test if EdU+, recently born cells are preferentially retained or lost. We therefore used a continuous, 48 hours long EdU pulse and studied if EdU+ fractions change in starving animals after 1, 2 or 5 days of chase (Fig. 3I). We indeed found that the EdU index significantly continuously increasing from 58.2±2.2% at 48hpf (G_1_: 71.9±1.1%) to 69.6±0.9% after 5 days of chase (G_1_: 73.4±0.4%)(Fig. S7C). Together, this increase suggests that cells incorporating EdU in the 48h after feeding tend to accumulate during the subsequent period of shrinkage and cell loss. However, as EdU signal intensity also decreases, we cannot exclude that the ∼10% increase is due to feeding-independent baseline proliferation. Future studies, for example by single-cell sequencing, are needed to further explore if cell loss affects specific cell populations or cell types.

Our study provides a benchmark for nutrient-dependent changes in body size, cell size, cell numbers and proliferation rates in *Nematostella vectensis*. The ease of inducing quiescence and cell cycle re-entry in a large fraction of cells by simple starvation and re-feeding makes *Nematostella* one of the most tractable and accessible animal systems to study the nutritional control of the cell cycle. The combination with the powerful genetic toolkit available (e.g., transgenesis, CRISPR/Cas9-mediated mutagenesis and knock-in) puts *Nematostella* in the spotlight as a powerful research organism to study the cellular and molecular basis of indeterminate growth control.

## Materials and Methods

### Animal culture

All *Nematostella* polyps were wildtype strains derived from the culture by Hand and Uhlinger (1992) and kept at 16ppm salt concentration (*Nematostella* medium) in a 25°C incubator without lights. After animals reached primary polyp stage, they were fed daily with mashed *Artemia salina* nauplii for the first week and with live nauplii thereafter. During feeding phases, *Nematostella* medium was replaced daily. Symbiotic and aposymbiotic clonal lines of Aiptasia strain CC7 (Grawunder et al., 2015), were maintained at 25 °C in 1,5l plastic boxes with filtered sea water (salinity approx. 32ppm). Symbiotic animals were kept at a 12:12 light cycle, while aposymbiotic animals were maintained in darkness and regularly screened for the absence of symbionts using fluorescent microscopy. Polyps were fed five days per week, with a full exchange of water twice a week, and a monthly scraping of the boxes. All experiments were performed on small polyps (∼3mm diameter) originating from pedal lacerates.

### Body size measurements in Nematostella vectensis and Exaiptasia pallidia

All experiments addressing growth and shrinkage were carried out in 6-well plates with individually housed animals except for experiment “140d starvation” in which pools of animals were kept in 3 replicate dishes. For shrinkage/starvation experiments, animals were fed until development of the secondary pair of full mesenteries (∼4-5 weeks after fertilization) as a morphological marker for juvenile polyps. Imaging was performed after relaxing animals using 0.1M MgCl_2_ in *Nematostella* medium for 20 minutes using a Leica M165 FC microscope. After masking tentacles, we measured body pixel area in ImageJ/Fiji (Schindelin et al., 2015) by detecting contrasting objects and automatically outlining polyp shape using “analyze particles”. We then converted pixel area to mm^2^ based on pixel size of a 2mm scale bar.

For body size measurements of Aiptasia, pedal lacerates of aposymbiotic and symbiotic animals of strain CC7 were left to regenerate into small polyps and fed until reaching a minimum size of ∼3mm diameter. 15-20 animals were selected for each of three replicate dishes per symbiosis condition and kept under 12:12h light cycle ((photon flux density of about 40 µmol m^−2^ s^−1^) in an incubator at 25°C. During the first 14 days of the assay (feeding phase), animals were fed daily for 1h with freshly hatched *Artemia* nauplii and washed 2-4 hours later to remove food waste. During the starvation period the assay dishes were cleaned, and sea water was replaced. Changes in body size were estimated by imaging and measuring of Aiptasia pedal disk area on anesthetized animals (Angeli et al., 2016). Polyps were relaxed for a minimum of 15min in 0.1 M MgCl_2_/sea water and pedal disks were imaged on an inverted microscope (Eclipse TE2000-S, Nikon). The images were imported into the ImageJ/Fiji software for analysis and pedal disk edge was outlined by hand using a polygonal selection tool. The resulting area was recorded and converted to mm^2^ using a scale bar.

### EdU labelling

Aiptasia or *Nematostella* polyps were relaxed for 15 min in 0.1M MgCl_2_ in sea water or *Nematostella* medium and transferred to sea water or *Nematostella* medium containing 300 µM EdU (5-ethynyl-2′-deoxyuridine, Invitrogen) 2%DMSO 0.1M MgCl_2_. Animals were incubated at RT for 60min for a short pulse and washed with 0.1M MgCl_2_ in sea water or *Nematostella* medium before being transferred to tissue homogenization tubes or fixed in 3.7% Formaldehyde in PBS overnight and stored in methanol at −20°C for microscopy sample preparation. During long EdU pulse and chase experiments, animals were kept in a 25°C incubator and no MgCl_2_ was added to the EdU medium, which was replaced every 24h.

Cells were permeabilized by incubation in 0.2% TritonX in 1% BSA/PBS for 15 min at RT and washed 1x with PBS (800g for 5 min, 4°C). The pellet was resuspended in 200 µl of freshly prepared Click-it reaction cocktail containing Alexa fluorophore azide (Invitrogen) and incubated for 30 min at RT in the dark. To reduce background, the cell suspension was washed twice with 1% BSA/PBS and stored at 4°C for analysis by flow cytometry within 48h. For whole mount samples, fixed polyps were washed twice in 1%BSA/PBS and permeabilized for 30 min at RT using 0.5% TritonX in 1% BSA/PBS. After three washes with 1%BSA/PBS the tissue was incubated in freshly prepared Click-it reaction cocktail (C10337, Invitrogen) with Alexa fluophore azide at RT for 30min in the dark, washed twice in 1%BSA/PBS and processed for cryosectioning.

### Cryosectioning

The sample polyps were left at 4°C to infiltrate with 25% sucrose, 20% OCT (Tissue-Tek, Sakura) in PBS overnight and frozen in 80% OCT/PBS on a metal block cooled by liquid Nitrogen. Using a Leica Cryostat, cross sections of 13μm were collected from the polyp midbodies and transferred to SuperFrost adhesion slides (Thermo Fisher). After 24h at RT, the tissue was re-fixed with 3.7% Formaldehyde/PBS for 10min and washed twice with PBS before mounting in 80% glycerol for confocal microscopy.

### Tissue homogenization and flow cytometer sample preparation

Depending on the experiment, 1 to 7 *Nematostella* polyps or 3 Aiptasia polyps constituted one biological sample and were transferred to tissue homogenization tubes (“C-tubes”, 130-096-334, Miltenyi Biotec) after they were relaxed. We added 5 ml of freshly prepared ACME solution (15% methanol, 10% glacial acetic acid and 10% glycerol in Milli-Q water, adapted from (García-Castro et al., 2021)) and incubated the samples for a total of 1h at RT with intermittent pulses of mechanical tissue disruption using the gentleMACS^TM^ tissue homogenizer (Miltenyi Biotec). The resulting homogenate was washed twice using 10 ml of ice-cold 1% BSA/PBS and pelleted for 5min at 800g (4°C). The cell pellet was gently resuspended in 1ml of 1% BSA/PBS and filtered through a pre-wetted 50 µm CellTrics strainer (Sysmex). For storage until all samples were collected, the cells were spun down (5min at 800g at 4°C) and taken up in 90% methanol to be kept at −20°C for up to a month. Before EdU labeling using a Click-It reaction and/or flow cytometry, cells were rehydrated by two washes with ice cold 1% BSA/PBS (5min at 400g, 4°C).

### Flow cytometry – estimating cell numbers, cell size and cell cycle distributions

Cell suspensions were stained with 1µg/ml FxCycle violet DNA dye (Invitrogen) on the day of the flow cytometry run. Flow cytometry was performed on a BD LSRFortessa (BD Life Sciences) and the resulting data was analyzed using FlowJoV10.8 (BD Life Sciences). For analysis of *Nematostella* whole body cell suspensions, we first excluded debris based on size and granularity in the FSC-A/SSC-A gate, with sub-gates based on FSC-A/FSC-H parameters and FSC-A/SSC-W parameters to remove potential cell doublets and high complexity events. We then gated particles based on DNA dye intensity in width over area and plotted a histogram of DNA dye in area on the linear scale to visualize the characteristic DNA dye intensity peaks expected from cells between 2N and 4N. These pre-selected events were considered the pool of cells from which analyses of cell size (median FSC-A intensity in arbitrary units), cell cycle composition (subfractions based on DNA signal intensity) and cell counting were performed. It was also used to record the fraction of EdU+ cells, by sub-gating on EdU labeling fluorescence in reference to negative controls. In the long term EdU pulse and chase experiments, the EdU+ population was further investigated using mean EdU fluorescence intensity (median value in arbitrary units) and cell cycle composition was established using DNA signal equivalent to the original gating. In cell counting experiments, 100 µl PBS containing a standardized quantity of 10^6^ 10µm FluoSpheres (F8834, Invitrogen) were added to 900 µl of sample homogenate of a single polyp to allow estimating the total cell number of cells per individual (see below). As these beads have comparable size to cells (see Fig.S6C) and we know their numbers, we can estimate the total number of cells per sample by counting beads using the following calculation:

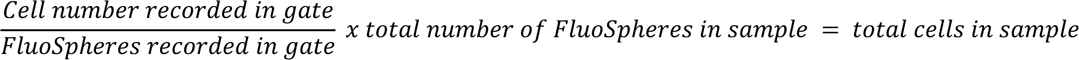

Proliferation assays in Aiptasia cell suspensions of both aposymbiotic and symbiotic animals were analysed with a slightly different strategy due to the presence of chlorophyllic autofluorescence in symbionts. After excluding events based on size and granularity (FSC-A/SSC-A, FSC-A/FSC-H, FSC-A/SSC-W) we gated on particles without symbiont autofluorescence / SSC-A and focused our analysis on this fraction. DNA signal intensity was again plotted as a histogram and events that fell in the expected range for cell cycle (2N-4N) were considered the reference population from which the fraction of EdU+ cells were recorded by comparison to negative controls.

### Cell counting to via Neubauer chamber

For validation of cell counts obtained by flow cytometry, cell suspension was transferred to a Millicell® Disposable Hemocytometer (MDH-2N1-50PK, Merck) counting chamber and according to manufacturer protocols, cells were manually counted and the mean number of cells was calculated.

### Confocal imaging

Polyp mid sections from EdU assays were imaged with a 20x oil-immersion lens using a Leica SP5 confocal microscope and standard PMT detectors. For validating cell size estimates, we stained the cell suspensions with Alexa Fluor™ 488 Conjugate Concanavalin-A (C11252, ThermoFisher) and Hoechst33342 and imaged individual cells on an Olympus FLUOVIEW FV3000 confocal microscope (standard PMT detectors) with a 60x oil-immersion lens. Images were processed and cropped in ImageJ/Fiji (Schindelin et al., 2012).

### Data analysis and mathematical modeling

Data visualization and statistical analysis were performed using R (R Development Core Team, 2023) with the libraries lmtest (Zeileis and Hothorn, 2002), ggplot2 (Wickham, 2016), gridExtra (Auguie, 2015), dplyr (Wickham et al., 2023), tidyverse (Wickham et al., 2019), ggridges (Wilke, 2022), survival (Grambsch and Therneau, 2000) and the Python libraries matplotlib (Hunter, 2007), Numpy (Harris et al., 2020), pandas (McKinney and others, 2010) and seaborn (Waskom, 2021). Detailed report on the models for analyzing growth and shrinkage data can be found in the Supplementary materials. In short, values for body size, cell number and cell size were natural logarithm transformed to achieve homoscedasticity and used as input linear regression models.

As especially during starvation, we anticipated several phases in the shrinkage response, we determined the intervals for the linear regression by employing a multi-phased changepoint-model approach that optimized fit by maximizing the log likelihood function. The optimization is done using the Nelder-Mead method for 1, 2 and 3 phases and using a simulated annealing algorithm for 4 or more phases. To estimate uncertainty in the parameter values, we employed bootstrap resampling (Berrar and Dubitzky, 2013). AIC (Akaike information criterion) values were calculated for model selection for each dataset. Positive slope values obtained from linear regressions of natural logarithm transformed data could then be interpreted as exponential growth (expressed as doubling time, T_d_) and negative slope values as exponential decay / shrinkage rates (expressed as halving time, T_1/2_) with an overlap of the slopes (± 2 s.e.m) considered it a similar response.

The multi-phased changepoint-model was mathematically defined as follows: for a number of *N* phases, we consider *N* − 1 changepoints. We assume that the first change-point is at day 0, denoted by *τ*_0_ = 0. We denote by *i* = 1, …, *N* the *i* th phase of the model, and we define a function *l*(*t*) that gives the phase corresponding to a given value *t*, that is, *l*(*t*) = max {*i*|*t* ≥ *τ_i_*_−1_ }. Then, this multi-phased, changepoint-model is a piecewise linear function defined by

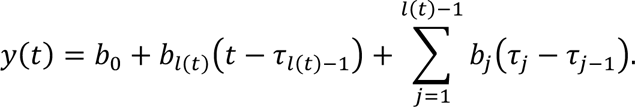

Equivalently, if we consider *i* = 1, …, *N* − 1 and we assume that *τ*_0_ = 0 and *τ_N_* = ∞, we can write the model by

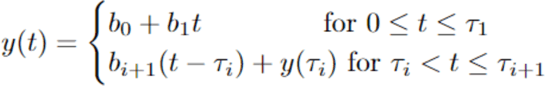

The space of parameters of the model is *θ* = (*b*_1_, *b*_0_, *σ*, *b*_2_, …, *b_N_*, *τ*_1_, …, *τ_N_*_−1_) where *σ* is the standard error of the normally distributed residuals, so that *σ* measures the spread of the observations from the line *y*(*t*). The log likelihood function *L*(*θ*|*y*(*t*)) of this model is then defined by

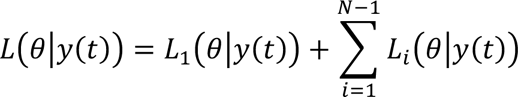

where

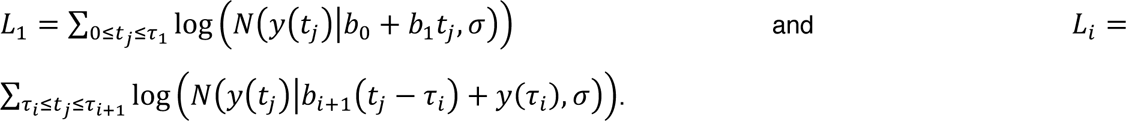

## Acknowledgements

We thank Frank Verhoek, Alena Pieters, Clément David for experimental support, Brith Bergum from the Flow cytometry Core Facility (University of Bergen) for technical guidance and support, Ira Mägele (Heidelberg University), Eilen Myrvold, Lavina Jubek and Brandon Mellin for help with the Aiptasia and *Nematostella* husbandry.

## Data availability

Data and code for the model fitting and statistics will be freely made available and accessible at <u>github.com/StochasticBiology/anemone-dynamics</u>.

## SUPPLEMENTARY FIGURES

**Fig. S1.**
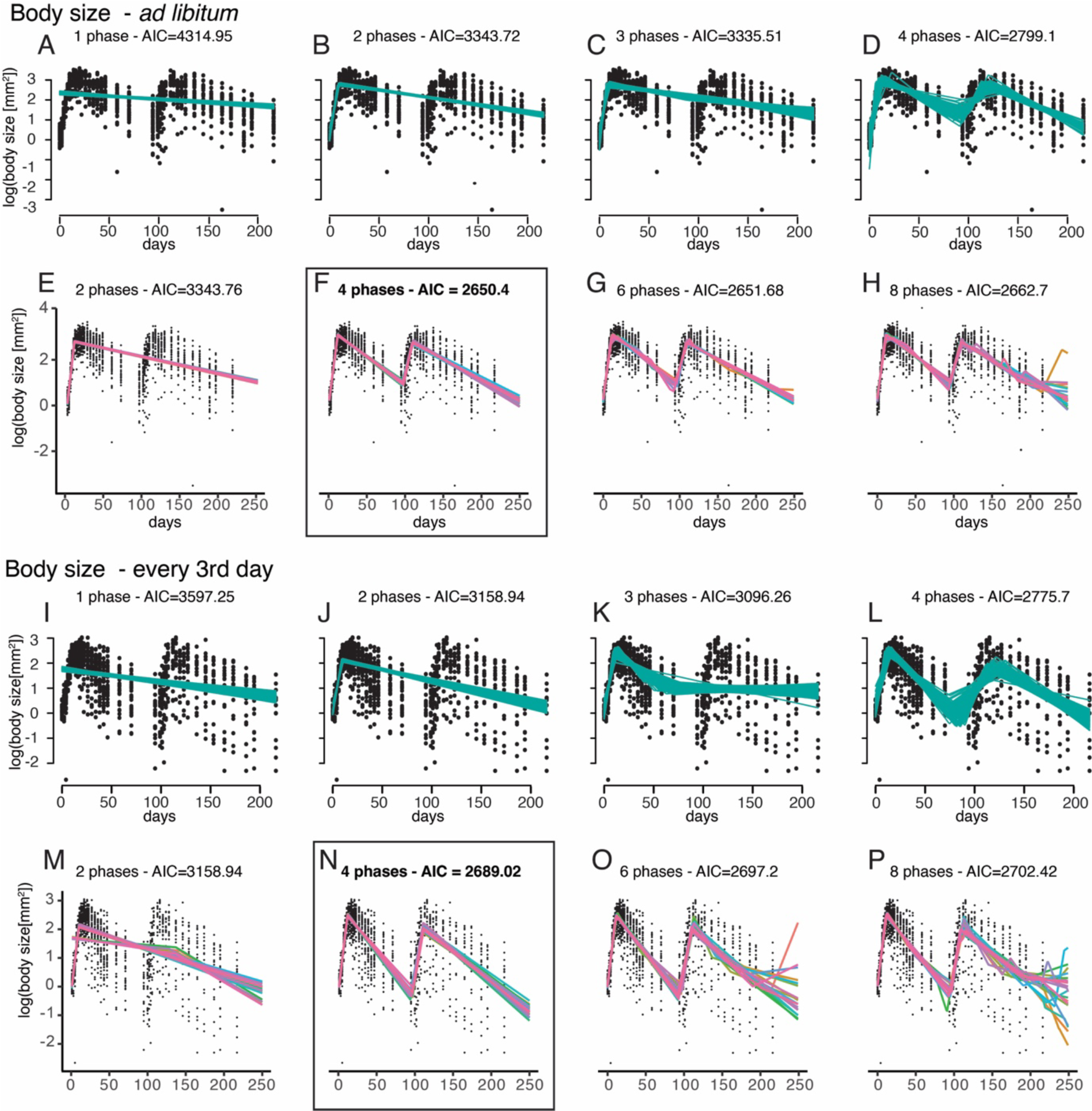
Selecting multi-phase linear models by maximizing log likelihood on log transformed, cyclic body size data. Comparison of model fits from maximising likelihood, with a Nelder-Mead fitting approach for 1-4 phases (*ad libitum* **A-D**, restricted feeding **I-L**) and a simulated annealing approach for 2-8 phases (*ad libitum* **E-H**, restricted feeding **M-P**), using the Akaike Information Criterion (AIC). The best statistical fit (lowest AIC) for both feeding conditions was a 4 phase model parameterised using simulated annealing (black boxes). For details on the model parameters see Supplementary Information.

**Fig. S2.**
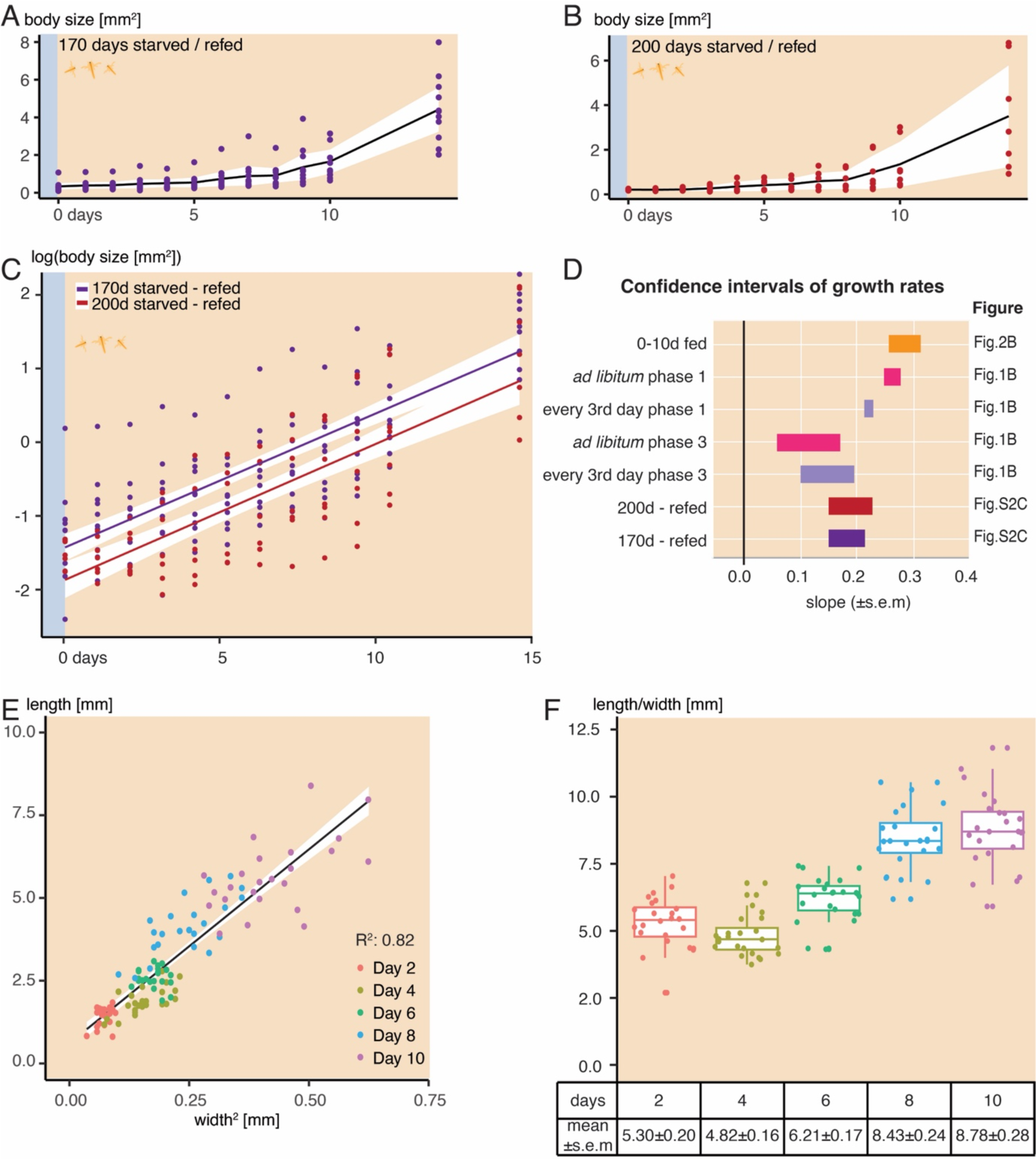
Stereotypy of growth during feeding or re-feeding in *Nematostella* juveniles. **(A-C)** Body size **(A,B)** and log-transformed body size **(C)** in *Nematostella* starved for 170 **(A)** or 200 days **(B)** after 14 days of *ad libitum* feeding. **(A, B)** show means and 95% confidence intervals connected across timepoints. **(C)** shows linear model on the log transformed values. **(D)** Estimates of growth rates from fitting linear models to log-transformed body size values. Note that slopes in animals fed *ad libitum* without prior starvation overlap, as well as the slopes from experiments with prior starvation (’refed’). **(E, F)** Length scales with the squared width in polyps fed *ad libitum* for 10 days. Shows means ±s.e.m. (see Fig. 2A; R^2^ = 0.82; N=24 per timepoint; see Supplementary Information: Geometry).

**Fig. S3.**
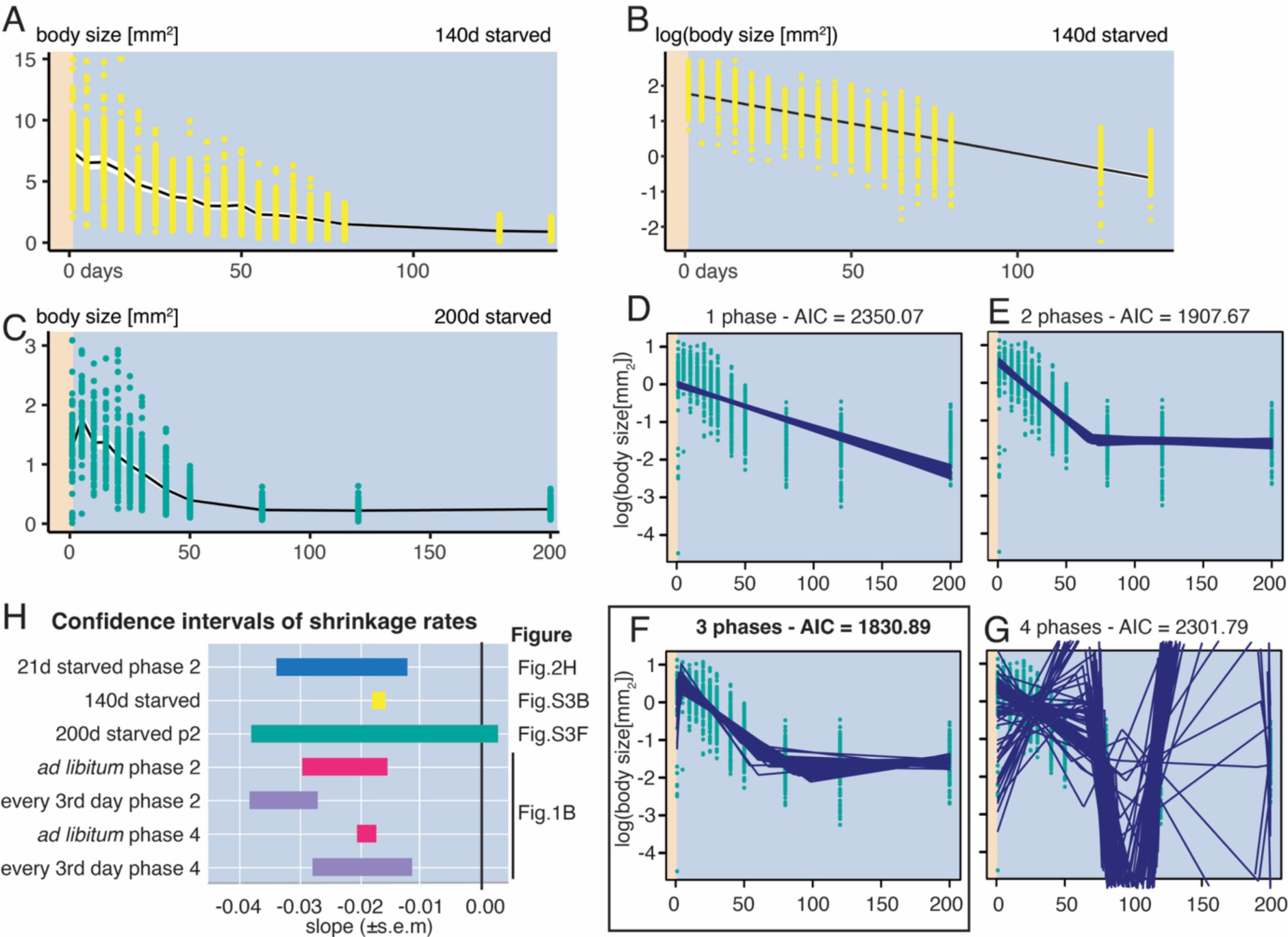
Stereotypy of shrinkage during starvation in *Nematostella* juveniles. **(A-C)** Body size **(A,C)** and log-transformed body size **(B)** during 140 **(A, B)** or 200 **(C-G)** days of starvation. **(A, C)** Mean values and 95% confidence interval were calculated per timepoint and connected (black line, white overlay). n=90. **(B)** shows linear regression and 95% confidence interval on natural logarithm transformed body size during 140 days of starvation (R^2^ = 0.53). **(C)** n=between 58 and 136 due to lethality during starvation. **(D-G)** A multi-phased linear regression model was fitted to the natural logarithm transformed values of body size during 200d starvation. Using a Nelder-Mead fitting approach, we defined a total of 3 phases (black box, for details on model selection see Supplementary Information) with 100 bootstraps visualized as blue lines. **(H)** Summary of estimated slopes (± 2 s.e.m.) for all experiments on *Nematostella* shrinkage. See see Supplementary Information for more details.

**Fig. S4.**
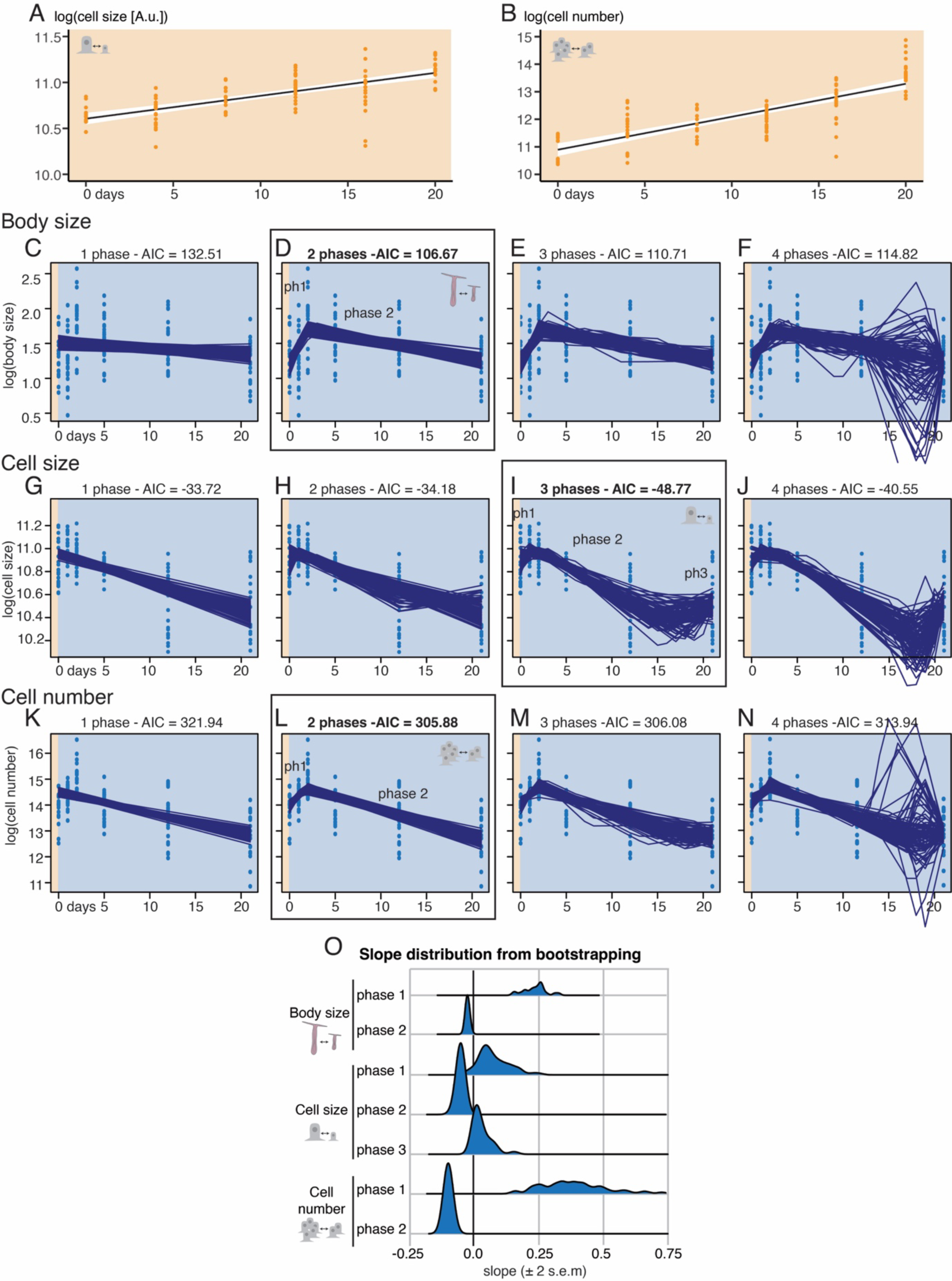
Body size, cell number and cell size dynamics during 10 days of feeding and 21 days of starvation in *Nematostella* juveniles. (A, B) Simple linear regression model on natural logarithm transformed values describe cell size **(A)** and cell number **(B)** changes during 10 days of *ad libitum* feeding. **(A)** Based on Fig.2D, R^2^ = 0.48, plotted: 95% CI. **(B)** Based on Fig.2C, R^2^ = 0.65 plotted: 95% CI. **(C-N)** Multi-phased linear regression models were fitted to the natural logarithm transformed values of body size **(C-F)**, cell size **(G-J)** and cell number **(K-N)** from individual polyps during 21 days of starvation. See also Fig.2G-J. 100 bootstraps (in blue lines) were plotted for a Nelder-Mead fitting approach to test 1 to 4 models and highlighted the best fit based on AIC (black boxes, see Supplementary Information). **(C-F)** A model with one growth and one shrinkage phase **(D)** had the best fit to describe body size changes over 21 days of starvation. **(G-J)** A 3 phase model **(I)** with an initial growth phase, a shrinkage phase and a plateau phase best described the dynamics of cell size changes. **(K-N)** A two phase model **(L)** with a post-feeding growth phase followed by a cell loss phase best described the dynamics of cell number changes. **(O)** Distribution of slope values (± 2 s.e.m.) obtained from bootstrapping for the individual phases in the chosen models for body size **(D)**, cell size **(I)** and cell number **(L)** changes.

**Fig. S5.**
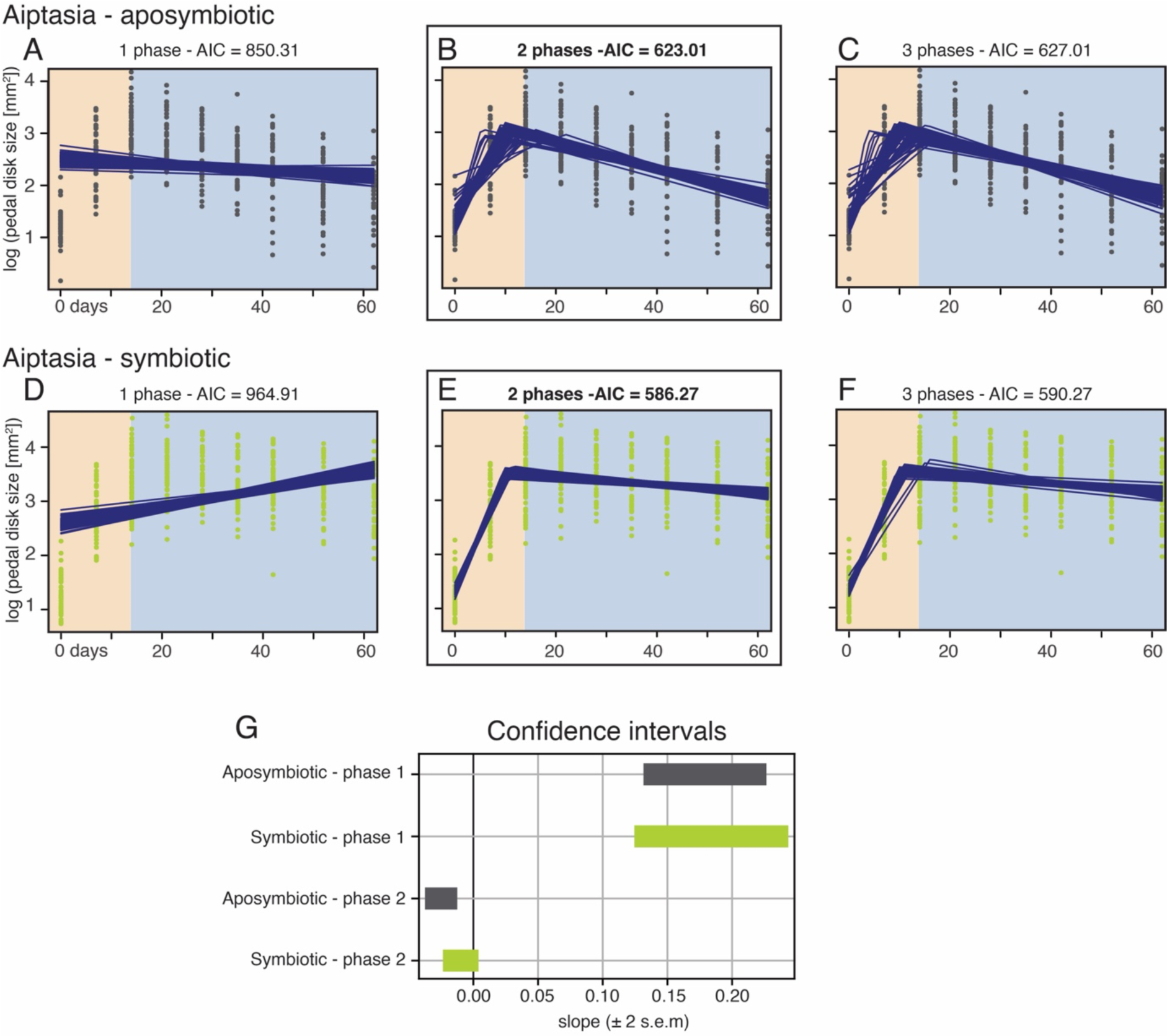
Aiptasia polyps show food related growth and shrinkage during starvation. **(A-F)** Multi-phased linear regression models were fitted to the natural logarithm transformed values of pedal disk area (Fig. 1H) in aposymbiotic **(A-C)** and symbiotic **(D-F)** Aiptasia polyps. For both conditions, two phases best described the dynamics of size changes with a growth phase followed by a shrinkage phase. See Supplementary Information for details on model selection. **(G)** Summary of slope values with 95% confidence interval for the best fitting change-point models overlap.

**Fig. S6.**
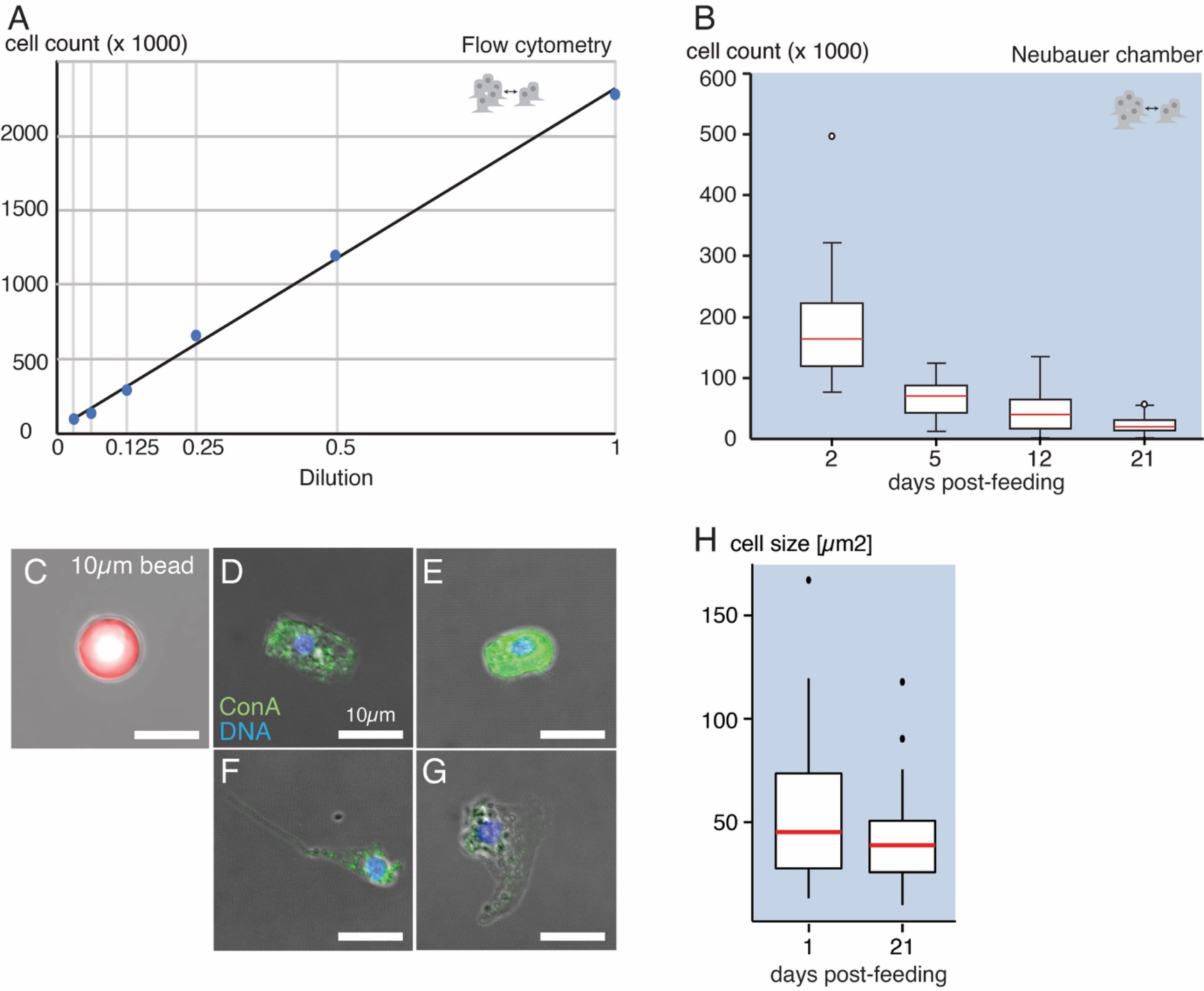
Validation of flow cytometry estimates for individual cell size and cell number. **(A)** Animal homogenate was diluted to benchmark scaling of cell counts obtained by flow cytometry. We found a near-perfect correlation (R² = 0,998) between the recorded cell count and the dilution coefficient. Average of 2 technical replicates shown. **(B)** Manual counting using a Neubauer chamber validates cell count experiments by flow cytometry. **(C-H)** Example images **(C-G)** and quantifications **(H)** of a 10 µm bead **(C)** or cells from cell suspensions **(D-G)** imaged by combined confocal and DIC microscopy to quantify average cell size. Green: Concanavalin A cytoplasmic stain (ConA). Blue: Hoechst33342 nuclear stain. **(C)** 10 µm beads used in the cell counting/flow cytometry experiment were confirmed to be of comparable size to the average cell (see Materials and Methods). **(H)** A trend towards smaller cells in 21d starved animals was observed, but the sample sizes (N_day 1_=42, N_day 21_=46) were likely too small and the range of their sizes likely too large to detect significant differences (pairwise t-test, p = 0.174).

**Fig. S7.**
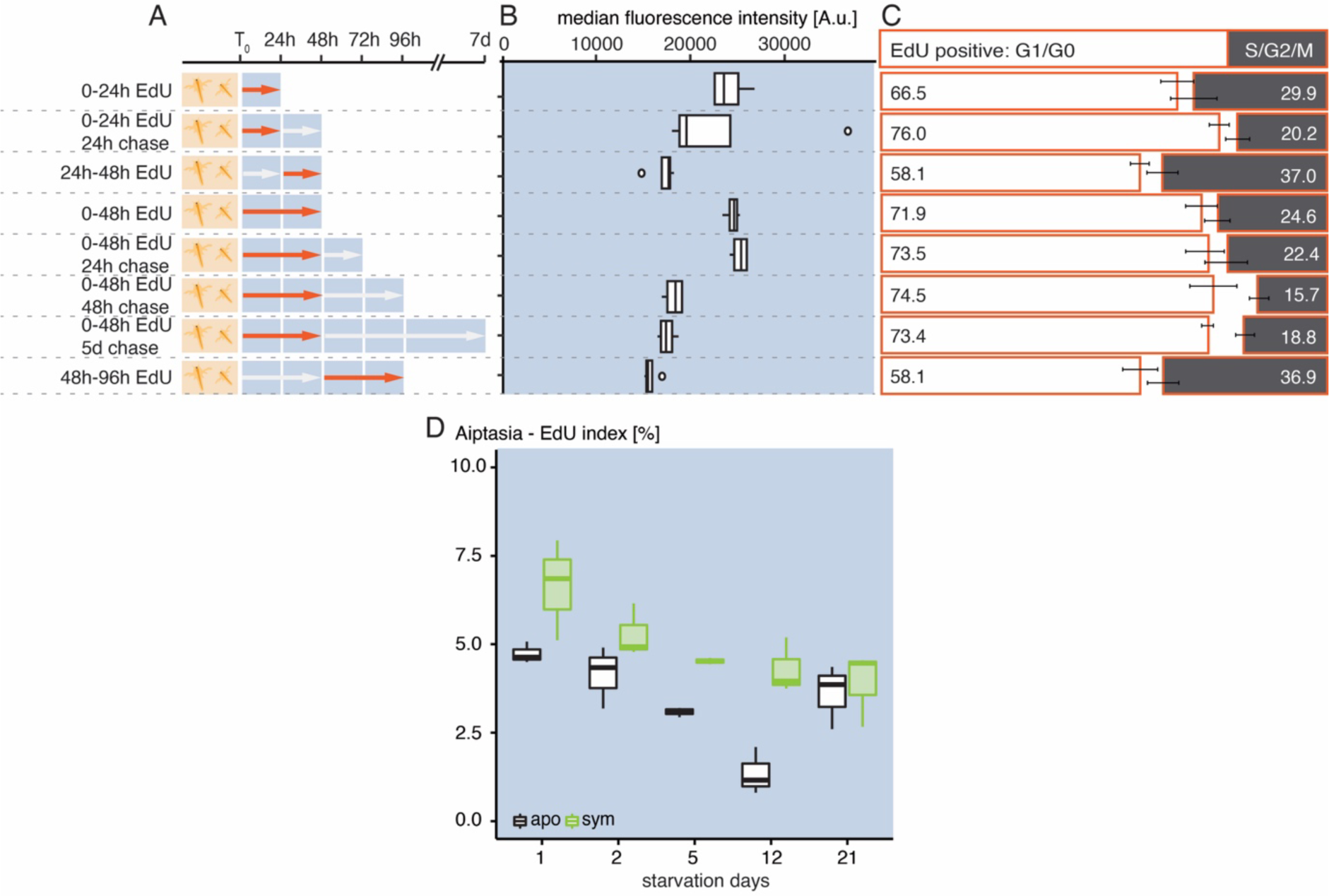
Starvation and feeding impact cell proliferation in *Nematostella* and Aiptasia. Experimental setup of EdU pulse-chase experiments **(A)**, median fluorescence intensity of EdU+ cells **(B)**, cell cycle distribution of EdU+ **(C)** and all polyp cells **(D)** in *Nematostella* juvenile (A-D) and EdU pulse labelling experiment in Aiptasia (E). **(B, C)** Same samples as in Fig. 3D. A.u.: Arbitrary units. **(C)** Cell cycle distribution of EdU+ cells as estimated by DNA signal intensity in flow cytometry. **(D)** Variations of the S-phase fraction in the same animals as in Fig. 2A. **(E)** 1h EdU pulse in Aiptasia strain CC7. n=3 per timepoint.

## Supplementary Information

### 1 Statistical models

In this section we present the statistical models used for the different data sets of body size changes and cellular changes in *Nematostella vectensis* in feeding and starvation periods. Estimates of parameter values are quantified with a 95% confidence.

#### 1.1 Software

Data preparation, statistical analysis and visualisations were performed using R [RStudio Team, 2020] and the R libraries lmtest [Zeileis and Hothorn, 2002], tidyverse [Wickham et al., 2019], ggplot2 [Waskom, 2021a], dplyr [Wickham et al., 2023], gridExtra [Auguie et al., 2017] and ggridges [Wilke, 2022]. We also used Python [Van Rossum and Drake Jr, 1995] and the Python libraries matplotlib [Hunter, 2007], numpy [Harris et al., 2020], pandas [pandas development team, 2020] and seaborn [Waskom, 2021b].

Data and code for the model fitting and statistics is freely available and can be accessed at GitHub github.com/StochasticBiology/anemone-dynamics.

#### 1.2 Simple linear regressions for feeding and refeeding periods

For a feeding period from day 0 to day 10, we have 144 data points for body size, 106 for cell number and 106 for cell size. We merge the data for these three measurements for each individual to get 106 data points, ignoring the unbalanced 38 data points from body size. This data is used for finding correlations between day and cell number, day and cell size, and body size and cell size and cell number. The original 144 data points for body size are used for correlations between day and body size.

The correlations found are of the form of simple linear regression models *y*(*x*) = *b*_1_*x*+*b*_0_ +*ɛ*, where *ɛ ∼ N*(0*, σ*) is noise that measures the spread of the observation *y* from the line *y*(*x*) = *b*_1_*x* + *b*_0_, *b*_1_ is the slope or coefficient and *b*_0_ the independent term or intercept. We usually work with log-transformed data. The parameter estimators and uncertainties of the models are shown in Table 1.

There are two main aspects that justify to work with the logarithmic transformation. First, we checked that it provides the property of homoscedasticity in the data, so that the variances of the observations *y_i_* for *i* = 1*, …, n* are all very similar at each discrete time point (day). This property is an essential assumption for regression analysis. Secondly, by using the logarithmic transformation, we can fit linear regressions and then naturally interpret these models as exponential growths or decays in the original variables (body size, cell number or cell size).

To study growth in refeeding periods, when animals are fed again after a period of starvation, we analyse data of animals that have gone trough starvation for 170 days, and animals that have gone trough starvation for 200 days. To explore whether the time over which animals have been starved makes a significant difference in how fast they grow when refeeded, we merged all data points from the previous two experiments, and we denote that merged data by ‘both’. The Akaike information criterion (AIC) value of the 170d model is 232.0, of the 200d model is 158.1, and of the merged model 408.1. Therefore, the AIC value of the model differentiating for how long animals were starved has a smaller AIC (232.0 + 158.1 = 390.1) than the model without differentiating starvation lengths (408.1). From here, we can select the first model and say that the length of the period of starvation matters to explain growth in the consecutive period of refeeding. However, in Table 1 we see that all the three slopes are practically identical, and the comparison of slopes via confidence intervals displayed in Fig S2D shows that the confidence interval for the 170d slope is almost entirely contained in the confidence interval for the 200d slope. Hence, we can select the second model, without differentiating the length of the previous starvation period, if we merely are interested in determining the growth rate.

**Table 1:** **Summary statistics of the simple linear regression models** in feeding and refeeding periods.

| Experiment | Slope | Intercept | $R^2$ |
| --- | --- | --- | --- |
| log body size 0-10d | $0.28 \pm 0.014$ | $-1.01 \pm 0.08$ | 0.91 |
| log cell number 0-10d | $0.239 \pm 0.0367$ | $10.89 \pm 0.23$ | 0.62 |
| log cell size 0-10d | $0.05 \pm 0.0099$ | $10.6 \pm 0.06$ | 0.49 |
| log body size refeeding 170d | $0.182 \pm 0.025$ | $-1.48 \pm 0.17$ | 0.61 |
| log body size refeeding 200d | $0.185 \pm 0.034$ | $-1.89 \pm 0.23$ | 0.59 |
| log body size refeeding both | $0.18 \pm 0.02$ | $-1.64 \pm 0.15$ | 0.57 |

We remark that for log cell number, the slope is *b*_1_ = 0.239 *±* 0.0367, and this corresponds to the rate of cell proliferation *λ* times the average proportion of cells that are either in the phase S, G2 or M, denoted by *S*, which is known from the data *S* = 0.1153. Then, the daily rate of cell proliferation is *λ ∼* 2.07.

##### 1.2.1 Correlations between body size and cell number and size from day 0 to day 10

Now we find a multiple linear regression to explore the correlations between body size and cell number and cell size. Let *b* denote the log transform of body size, *n* the log transform of cell number and *s* the log transform of cell size. All these variables are dependent on time in days *x*, but we consider how they scale together over the measured range of times. The parameter estimators of this multiple regression are summarised in Table 2 below.

**Table 2:** **Summary statistics of the multiple regression model** for log body size *b* as a function of log cell size *s* and log cell number *n*.

| Experiment | Slope for $s$ | Slope for $n$ | Intercept | $R^2$ |
| --- | --- | --- | --- | --- |
| $b \sim s + n$ | $1.04 \pm 0.66$ | $0.63 \pm 0.15$ | $-18.5 \pm 6.0$ | 0.68 |

We also explore the effect that cell number and cell size independently have on body size. For that, we fit two simple linear regressions whose parameter estimates are summarised in Table 3 below.

**Table 3:** **Summary statistics of the two simple regression models** for log body size *b* as a function of log cell size *s* and log cell number *n*, respectively.

| Experiment | Slope | Intercept | $R^2$ |
| --- | --- | --- | --- |
| $b \sim s$ | $2.9 \pm 0.6$ | $-31.6 \pm 6.5$ | 0.48 |
| $b \sim n$ | $0.8 \pm 0.1$ | $-9.3 \pm 1.4$ | 0.65 |

#### 1.3 Multi-phase models for starvation periods and cyclic periods

The changes in body size and at a cellular level are more complex to understand during starvation periods. This is because the animals undergo different phases of growth and degrowth when there is a prolonged lack of food. Therefore, simple regression models do not longer capture the biological processes as accurately as they do for body growth and cell growth and proliferation in feeding periods. We approach this problem by defining multi-phase linear models whose parameter values 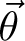 are inferred by maximising their log likelihood function 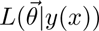 for given data *y*(*x*). We maximise the log likelihood function using the function *optim* in R that employs the Nelder-Mead method (when we want to explore one, two or three phases), or using a custom-written simulated annealing method in C++ (when we want to explore four or more phases). To obtain uncertainties for the predictors and parameter estimates, we employ bootstrap resampling [Berrar and Dubitzky, 2013] 300 times for each dataset, to obtain a distribution of estimates for the slopes and changepoints. From the distributions, we obtain 95% confidence intervals as the values between the 2.5*^th^* and the 97.5*^th^* percentiles in the distribution. The model selection is done by choosing the model with the number of phases that has lowest AIC value. Therefore, we select a 2-phase model for the logarithmic transform of body size Fig S4D and for the logarithmic transform of cell number Fig S4L, a 3-phase model for the logarithmic transform of cell size Fig S4I and a 4-phase model for the logarithmic transform of body size in cyclic periods of feeding, starvation, refeeding and restarvation for *ad libitum* conditions (AL) Fig S1F and restricted conditions every 3 days of feeding (RES) Fig S1N.

A multi-phase model for a number of *N* phases has *N −* 1 changing-time-points. We impose that the first changing-time-point is *τ*_0_ = 0. We denote by *i* = 1*, …, N* the *i*th phase of the model, and we define a function *l*(*t*) that gives the phase corresponding to a given value *t*, that is, *l*(*t*) = max*{i | t ≥ τ_i−_*_1_*}*. Then, this model is a piecewise linear function defined by:

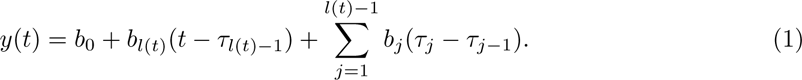

Equivalently, if we consider *i* = 1 *…, N −* 1 and we assume that *τ*_0_ = 0 and *τ_N_* = *∞*, we can write the multi-phase, piecewise linear model by:

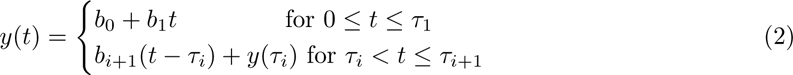

The set of parameters of the model is 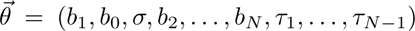 where *σ* is the standard error of the normally distributed residuals, so that *σ* measures the spread of the observations from the line *y*(*t*). The log likelihood function 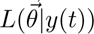 of this model is then defined by:

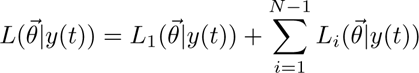

where

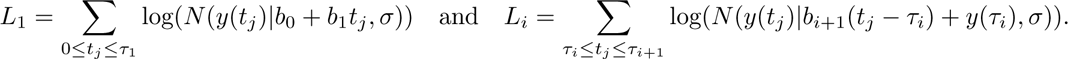

The general structure in these multi-phase models is a first phase of growth, with positive slope, a second phase of degrowth, with negative slope, and a third phase of growth in the case of cell size, as cells reduce their size until a threshold where they cannot become smaller, and from there they become bigger again. The models for cyclic periods AL and RES have a a first phase of growth, a second of degrowth, a third of growth and a fourth of degrowth.

To clarify notation, We have two datasets for body size measurements in long starvation periods. We denote by (*i*) the data for around 170 days and by (*ii*) the data for 200 days. The model for the long starvation (*i*) is a simple linear regression model, as the data has a good structure for a single phase and this one-phase model gives a high coefficient of determination *R*^2^ of 0.53, as seen in Table 4 below.

**Table 4:** **Summary statistics of the multi-phase linear models** for starvation periods and cyclic periods. Note that the row log body size long (i) is a simple linear regression model.

| Experiment | First slope | Second slope | Third slope | Fourth slope | $R^2$ |
| --- | --- | --- | --- | --- | --- |
| log body size 0-21 | $0.23 \pm 0.08$ | $-0.02 \pm 0.01$ | - | - | 0.20 |
| log cell number 0-21d | $0.44 \pm 0.02$ | $-0.10 \pm 0.02$ | - | - | 0.43 |
| log cell size 0-21d | $0.09 \pm 0.11$ | $-0.049 \pm 0.001$ | $0.06 \pm 0.07$ | - | 0.45 |
| log body size long (i) | $-0.017 \pm 0.0004$ | - | - | - | 0.53 |
| log body size long (ii) | $-0.004 \pm 0.008$ | $-0.01 \pm 0.02$ | $0.002 \pm 0.013$ | - | 0.52 |
| log body size AL | $0.25 \pm 0.02$ | $-0.025 \pm 0.002$ | $0.138 \pm 0.025$ | $-0.018 \pm 0.001$ | 0.55 |
| log body size RES | $0.22 \pm 0.01$ | $-0.033 \pm 0.004$ | $0.15 \pm 0.03$ | $-0.021 \pm 0.004$ | 0.28 |

We remark that for log cell number, the cell proliferation rate in the first phase of growth is *λ ∼* 2.5, computed as for the feeding period *b*_1_ = *λS*, since here the average cellular proportion in either S, G2 or M phase is *S* = 0.1633 and the slope is *b*_1_ = 0.44 *±* 0.024.

We report changepoints *τ* by finding their 95% confidence intervals in an analogous way as for the slopes. These are:

**Table 5:** **Confidence intervals of the time changepoints of multi-phase linear models**. 95% confidence.

| Experiment | $\tau_1$ | $\tau_2$ | $\tau_3$ |
| --- | --- | --- | --- |
| log body size 0-21 | [1.9, 2.6] | - | - |
| log cell number 0-21d | [1.1, 2.0] | - | - |
| log cell size 0-21d | [0.8, 6.0] | [9.9, 19.2] | - |
| log body size long (ii) | [0.1, 5.9] | [47.2, 97.3] | - |
| log body size AL | [7.9, 15.3] | [3.1, 102.7] | [112.5, 134.1] |
| log body size RES | [10.6, 16.6] | [62.7, 92.2] | [110.7, 136.2] |

##### 1.3.1 Correlations between body size and cell number and size from day 0 to day 21

We explore how cell number and cell size influence body size in each growth and degrowth phase during starvation. To do this, we see in the plots of the selected models that the changepoint is roughly day 2, so we fit a multiple linear regression for the data between day 0 and day 2, both included, for growth and between day 2 and day 21, both included, for degrowth.

Following the notation as for the correlations in feeding periods, we have the summary statistics Table 6 and Table 7 below.

**Table 6:** **Summary statistics of the multiple regression model** for log body size *b* as a function of log cell size *s* and log cell number *n*.

| Experiment | Slope for $s$ | Slope for $n$ | Intercept | $R^2$ |
| --- | --- | --- | --- | --- |
| 0-2d: $b \sim s + n$ | $0.33 \pm 0.48$ | $0.41 \pm 0.12$ | $-8.1 \pm 5.7$ | 0.4 |
| 2-21d: $b \sim s + n$ | $-0.01 \pm 0.34$ | $0.20 \pm 0.09$ | $-1.0 \pm 2.9$ | 0.3 |

**Table 7:** **Summary statistics of the two simple regression models** for log body size *b* as a function of log cell size *s* and log cell number *n*, respectively.

| Experiment | Slope | Intercept | $R^2$ |
| --- | --- | --- | --- |
| 0-2d: $b \sim s$ | $0.13 \pm 0.60$ | $-0.04 \pm 6.72$ | 0.0028 |
| 0-2d: $b \sim n$ | $0.4 \pm 0.118$ | $-4.3 \pm 1.7$ | 0.39 |
| 2-21d: $b \sim s$ | $0.2 \pm 0.3$ | $-4.0 \pm 2.7$ | 0.15 |
| 2-21d: $b \sim n$ | $0.20 \pm 0.06$ | $-1.2 \pm 0.8$ | 0.30 |

We notice that in both growth and degrowth phases during starvation periods, we have no confidence to say that cell size influences body size, as log cell size can be scaled by 0 (slope for *s* in Tables 6 and 7). We find that in the early phase of growth, cell number explains almost 40% of the variability of body size. In the latest phase of degrowth, cell number explains 30% of the variability of body size (Table 7).

#### 1.4 Comparison of growth and degrowth rates across experiments

We say that growth and degrowth rates, corresponding to the slopes of the models for each experiment, coincide if their confidence intervals intersect. This can directly be seen in the overlap of the confidence interval bars Figs 1B, S2D and S3H. We say that growth and degrowth rates are significantly different between the experiments if their confidence intervals do not intersect. In this paper we always work with 95% confidence.

Overall most growth rates coincide, suggesting a certain level of universality in how fast animals increase their body size when they are fed (or refed). Significant differences are found between the growth rate in 0-10d and each growth rate in feeding RES, refeeding RES and AL and refeeding after 170d and 200d. We also see that animals experiencing refeeding in AL and in RES grow at lower rates than animals experiencing refeeding after 170 or 200 days. These higher rates of refeeding after 170 and 200 days slightly match with the growth rate for feeding in RES.

All degrowth rates match with each other for the different datasets, except for the rate of the first degrowth phase in RES and the rate of long starvation (i), and the rate of the second degrowth in AL and the first degrowth phase in RES. These big similarities in how animals shrink across different biological conditions might be pointing to a lack of habituation in starvation. Another important result of this analysis is that for the starvation period between 0 and 21 days, the first phase of growth lasting approximately until day 2, happens with at a rate very similar to the rates in feeding 0-10d, feeding AL, feeding RES, refeeding 170d and refeeding 200d.

We can conclude that growth rates are one order of magnitude higher than shrinkage rates, so animals grow around ten times faster than they shrink.

### 2 Geometry

From the correlation between body size and cell number and cell size in growth during feeding periods, Table 2, we can give insights in how cell proliferation and cell growth increase the body size by studying the geometry of the body.

*Nematostella vectensis* has a cylindrical body shape, its body size is measured as the area (in *mm*^2^) of a rectangular projection of this cylinder into a viewing plane. For *r* denoting the radius of the base of the cylinder and *h* the height, the formulas of areas and volumes that we need are the following: area of a rectangle (body size)

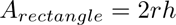

which can change in width 2*r* or in height *h*. Area of a cell (cell size)

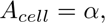

volume of a cylinder:

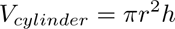

which can change radially *πr*^2^ or in elongation *h*, and lastly, volume of a cube with edge length *l*; *V_cube_* = *l*^3^.

The parameters of the correlation of body size and cell number and cell size during growth in feeding periods are in Table 2. The slope for the variable log cell number is 0.63 *±* 0.155, which is approximately 2*/*3, and the slope for the variable log cell size is 1.04 *±* 0.66, which can be rounded to 1. These values appear in geometric formulas, so we look closely how to connect the geometries of the body and cells of *Nematostella vectensis* to known scaling laws. First, we rewrite the correlation equation in Table 2 with the approximations presented and taking the inverse of the logarithmic transformation:

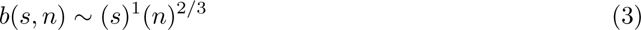

so that here *b, n* and *s* are denoting body size, cell number and cell size, respectively.

As cell size, like body size, is measured as a 3D volume projected into an observed 2D plane, the linear scaling of observed body size with observed cell size intuitively corresponds to body size scaling like cell size. The 2*/*3 scaling with cell number suggests that increasing cell number may contribute sublinearly to either the length or radius, or both, of the 3D body form, which is unsurprising given the complexity of the body plan. Further detailed physiological work will dissect this relationship further.

### 3 Doubing times, half-lives and interval loss rates in starvation

*Nematostella vectensis* undergo exponential growth and exponential decay as degrowth, of the form *N*(*t*) = *N*_0_*e^rt^* where *r* is the growth rate (if positive) or degrowth rate (if negative), corresponding to the slope of the model. In this scenario, it is informative to compute the doubling times *T_d_* and the half-lives *t*_1_*_/_*_2_ to know the time that it takes for the initial quantity to double in number or to reduce to half, respectively. The doubling time *T_d_* formula is

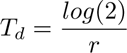

where *log* is the natural logarithm in base *e*. The half-life *t*_1_*_/_*_2_ formula is:

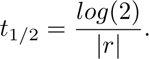

**Table 8:**
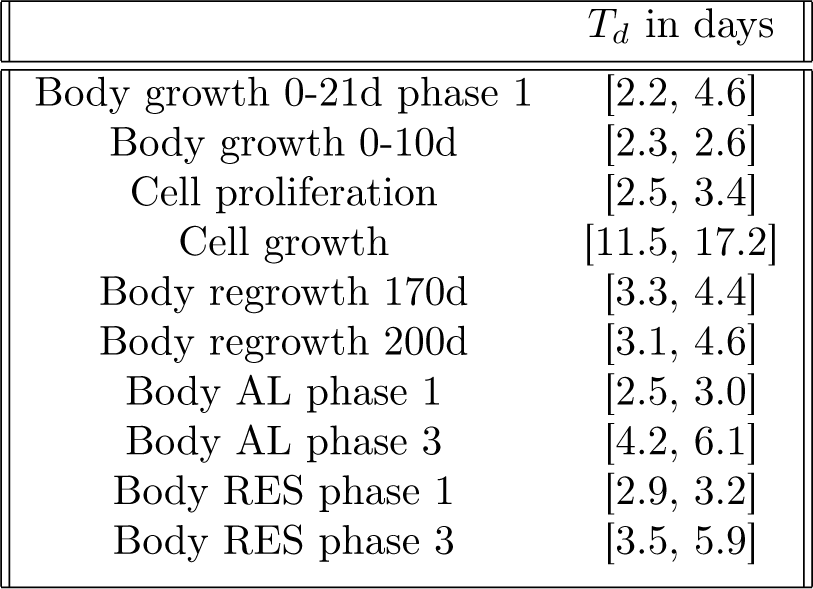
**Doubling times** for exponential growth models. 95% confidence intervals.

**Table 9:**
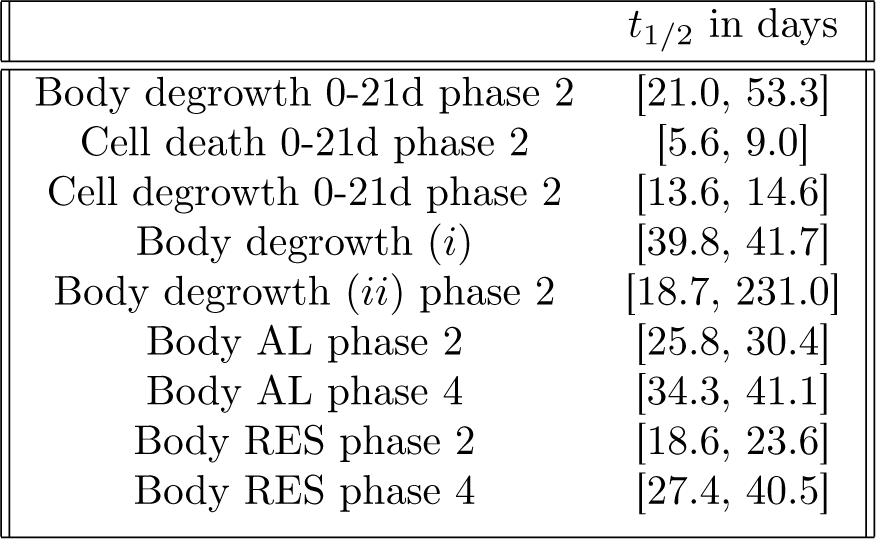
**Half-lives** for exponential decay models. 95% confidence intervals.

*Nematostella vectensis* doubles their body size in approximately between 2 and 4 days, their cell number doubles in between 2 and 3 days and their cell size doubles in between 11 and 17 days. These animals halve their body size in between 20 and 50 days, although if the animals start the starvation period with smaller bodies (as it is the case in RES compared to AL), it takes considerably less days to halve their size. This might indicate that bigger individuals are more resilient in starvation periods than smaller ones, although for second phases of starvation (phase 4 in AL and RES) there is almost no difference.

Next we display loss rates for each specific time interval in Table 10 below. These rates correspond to the slopes of simple linear regressions that have been fitted for each time interval. We remark that these loss rates are equal to the absolute value of the slope mean estimates of the regressions, where the slopes are negative as we are in degrowth phases.

**Table 10:** **Loss rates** for each time interval.

| Time interval | Body size | Cell number | Cell size |
| --- | --- | --- | --- |
| 2-5d | 0.085 | 0.325 | 0.033 |
| 5-12d | 0.0124 | 0.066 | 0.046 |
| 12-21d | 0.0261 | 0.073 | 0.00027 |

We remark that these results point towards cell size reaching a threshold at around day 12 of starvation, so that it does not shrink further and maintains its size almost constant.

### 4 Additional models for Aiptasia

We have two datasets of the logarithmic transformation of body size measurements with respect to time in days for the species Aiptasia. One dataset is for individuals that do not contain endosymbionts, called ‘apo’, and the other for individuals that do contain symbionts, called ‘symb’. The data is cyclic, meaning that animals go through a period of feeding followed by a period of starvation. Therefore, the model approach is the multi-phase model presented before for starvation and cyclic periods in *Nematostella vectensis*. We fit a simple linear regression (1-phase model), a 2-phase model and a 3-phase model, and we see that the 2-phase model gives lowest AIC value for both ‘apo’ and ‘symb’ Fig S5B,E. Hence, we select 2-phase models to describe and predict the changes in body size in Aiptasia when fed and consecutively starved, having a first phase of growth and a second of degrowth.

We next compare the slopes (corresponding to growth and degrowth rates) between ‘apo’ and ‘symb’ individuals, and we can conclude that both growth rates are almost identical between them, with ‘sym’ taking some higher values Fig S5G. This is also seen in their doubling times Table 4. So animals either with or without symbionts grow almost equally fast when fed, but those with symbionts can grow slightly faster. For degrowth rates, corresponding to the second phase of the models, individuals with symbionts can shrink slower than those without symbionts Fig S5G. This is also seen in the values of their half-lives Table 4.

**Table 11:**
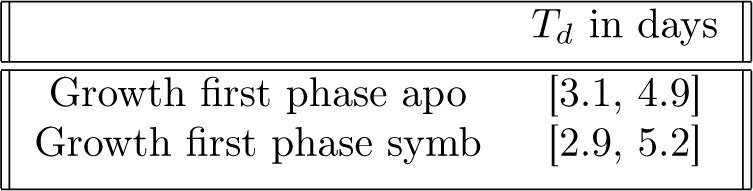
**Doubling times** for Aiptasia. 95% confidence intervals.

**Table 12:**
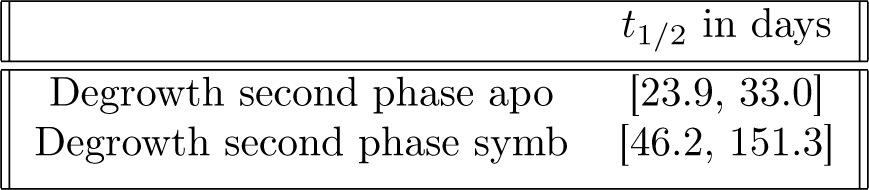
**Half-lives** for Aiptasia.95% confidence intervals.

**Table 13:** Confidence intervals of the changepoint of 2-phase linear models.

| Experiment | $\tau_1$ |
| --- | --- |
| apo | [4.96, 17.38] |
| symb | [9.42, 11.82] |

